# Improving Functional Muscle Regeneration in Volumetric Muscle Loss Injuries by Shifting the Balance of Inflammatory and Pro-Resolving Lipid Mediators

**DOI:** 10.1101/2024.09.06.611741

**Authors:** Thomas C. Turner, Frank S. Pittman, Hongmanlin Zhang, Lauren A. Hymel, Tianyi Zheng, Monica Behara, Shannon E. Anderson, Julia Andraca Harrer, Kaitlyn A. Link, Mashoor Al Ahammed, Kristal Maner-Smith, Xueyun Liu, Xuanzhi Yin, Hong S. Lim, Matthew Spite, Peng Qiu, Andrés J. García, Luke J. Mortensen, Young C. Jang, Nick J. Willett, Edward A. Botchwey

## Abstract

Severe tissue loss resulting from extremity trauma, such as volumetric muscle loss (VML), poses significant clinical challenges for both general and military populations. VML disrupts the endogenous tissue repair mechanisms, resulting in acute and unresolved chronic inflammation and immune cell presence, impaired muscle healing, scar tissue formation, persistent pain, and permanent functional deficits. The aberrant healing response is preceded by acute inflammation and immune cell infiltration which does not resolve. We analyzed the biosynthesis of inflammatory and specialized pro-resolving lipid mediators (SPMs) after VML injury in two different models; muscle with critical-sized defects had a decreased capacity to biosynthesize SPMs, leading to dysregulated and persistent inflammation. We developed a modular poly(ethylene glycol)-maleimide hydrogel platform to locally release a stable isomer of Resolvin D1 (AT-RvD1) and promote endogenous pathways of inflammation resolution in the two muscle models. The local delivery of AT-RvD1 enhanced muscle regeneration, improved muscle function, and reduced pain sensitivity after VML by promoting molecular and cellular resolution of inflammation. These findings provide new insights into the pathogenesis of VML and establish a pro-resolving hydrogel therapeutic as a promising strategy for promoting functional muscle regeneration after traumatic injury.

## 1. INTRODUCTION

Volumetric muscle loss (VML) is a loss of muscle mass that fails to regenerate without intervention and usually occurs in traumatic injuries, such as motor vehicle accidents, military combat, or surgical resection^1^. The current gold standard treatment for VML is surgical reconstruction with muscle flap autografts or free tissue transfer. However, this treatment is often complicated by excessive fibrosis and fatty infiltration, which can impair regeneration and lead to significant long-term disabilities and chronic pain^2^. Inflammation is a key component of VML pathology and creates a significant challenge in treating VML. Uncontrolled or persistent inflammation contributes to the initial muscle damage, can impede regeneration, and exacerbate functional strength deficits and chronic pain^3^. Therefore, understanding the mechanisms of inflammation resolution and identifying therapeutic targets to promote that resolution is critical to improving outcomes for VML patients.

As with other inflammatory tissue disorders such as osteoarthritis, periodontitis, and diet-induced obesity, there is strong interest in developing therapeutics that modulate inflammation to enhance tissue regeneration, improve harmful symptoms, or cure disease^4–6^. The current approach to treating excessive inflammation and the resulting collateral damage is largely based on blocking or inhibiting pro-inflammatory chemical mediators to suppress the inflammatory cascade^7,8^.

Whereas pro-inflammatory eicosanoids such as Prostaglandins and Leukotrienes are essential in initiating the first stages of inflammation, excessive immune response can lead to impaired regeneration and chronic pain^9^. However, inhibiting key pathways contributing to excessive inflammation can also lead to immune suppression, poor wound healing, damage to collateral tissues, and other unwanted side effects^10,11^.

Several families of specialized pro-resolving lipid mediators (SPMs) have recently been identified, including Lipoxins, Resolvins, Protectins, and Maresins; these SPMs act through distinct G protein-coupled receptors (GPCRs) that regulate the successful resolution of inflammation^12,13^. Lipidomic analyses of human tissues in various inflammation and tissue regeneration models have shown the potential role and importance of SPM pathways in tissue pathologies and recovery^14,15^. In pre-clinical murine models of muscle injury and regeneration, the concentration of pro-inflammatory lipid mediators and SPMs are correlated with periods of inflammation and resolution/regeneration, respectively. These studies have characterized the lipid mediator response to acute injury and its role in mediating wound healing^16–19^. However, the lipid mediator response following a non-healing VML muscle injury has yet to be comprehensively characterized.

Therapeutic delivery of SPMs through systemic or topical administration has been shown to promote wound healing, reduce pain, and mitigate fibrosis^20–22^. Systemic administration of exogenous lipid mediators, namely D-series Resolvins RvD1 and RvD2, significantly improved muscle regeneration in both acute injury models and Duchenne muscular dystrophy^17–19,23^. D-series Resolvins increase muscle regeneration through both actions on immune cells promoting resolution as well as direct signaling on progenitor cells to increase proliferation^19^. However, endogenous SPMs are highly prone to metabolic inactivation *in vivo*, which often limits their effects with traditional delivery methods^24,25^. Furthermore, SPMs carry out their functions via short-range signaling interactions in a localized and targeted manner^14^. Thus, the development of biomaterial platforms that protect these lipids from enzymatic inactivation and facilitate their local, short-range interactions is an emerging and impactful innovation in resolution pharmacology, especially in the context of traumatic injuries.

The objective of this study was to: 1) investigate the temporal shifts in pro-inflammatory and pro-resolving lipid mediator response after critically-sized VML injuries at various time points (up to 2 weeks post-VML), 2) engineer a novel hydrogel biomaterial platform to provide local delivery of AT-RvD1. We hypothesized that local delivery of AT-RvD1 would shift the lipid mediator response to resolve chronic inflammation and promote functional muscle regeneration after critically-sized VML injury. Using VML models in both the quadriceps and spinotrapezius, this hypothesis was rigorously assessed using comprehensive Liquid Chromatography-Mass Spectrometry (LC-MS) of lipid mediator biosynthesis, flow cytometry of immune cell dynamics, hindlimb functional and pain outcome metrics, and structural and cellular muscle regeneration outcomes. The local and early delivery of AT-RvD1 via synthetic hydrogels significantly altered *in vivo* biosynthesis of SPMs, initiated a pro-regenerative shift in the immune cell composition and muscle progenitor cells, decreased limb pain sensitivity and enhanced functional muscle regeneration to a level indistinguishable from healthy muscle.

## 2. MATERIALS AND METHODS

### 2.1 Hydrogel synthesis

Four-arm PEG macromer (10-kDa molecular weight) end functionalized with maleimide (>95% purity; Laysan Bio) at a final density of 4.5% (w/v) was used for all hydrogel formulations as described previously^26,27^. PEG macromers were functionalized with RGD peptide (GRGDSPC), cross-linked with the protease-degradable cysteine-flanked peptide VPM (GCRDVPMSMRGGDRCG) (AAPPTec) in 0.5 M MES buffer (pH 5.5). The final concentration of RGD was 1.0 mM. The cross-linker was adjusted to consume non-reacted maleimide groups remaining on PEG macromers. Gels were also loaded with 100 ng of AT-RvD1 (4 μg/mL) (Cayman Chemical). For hydrogels used in animal studies, all components were filtered through a spin column after pH measurements and kept under sterile conditions until injection into the animals and *in situ* polymerization.

### 2.2 Quadricep VML surgery and hydrogel implantation

All animal procedures were conducted according to protocols approved by Georgia Institute of Technology Institutional Animal Care and Use Committee. Male C57BL/6J mice (8-12 weeks old, Jackson Laboratory) were used for all animal studies. Surgical procedure was performed as previously reported^28^. Briefly, the left hindlimb was prepped and sterilized. An incision was made above the quadriceps and either a 2 mm (for sub-critical VML analysis) or a 3 mm (for critical VML analysis) biopsy punch (VWR, 21909-136) was used to make a full-thickness muscle defect^28^. Hydrogel components were mixed and immediately pipetted into the defect space for *in situ* cross-linking. The skin was closed, and animals were allowed to heal for specified number of days before euthanasia by CO_2_ inhalation. For systemic administration of AT-RvD1, immediately after VML surgeries were performed and the skin was closed, the animals were treated with 100 ng AT-RvD1 via once intraperitoneal injection.

### 2.3 Spinotrapezius VML surgery and hydrogel implantation

A 2 mm-diameter full thickness defect in the spinotrapezius muscle was created as previously described^29^. Briefly, a longitudinal 1-inch incision was made just after the bony prominence of the shoulder blade. The overlying fascia was dissected away and the spinotrapezius muscle was identified. The edge of the spinotrapezius was reflected and positioned against a sterile piece of wood and a full-thickness defect was made through the muscle using a 2 mm biopsy punch. Hydrogel components were mixed, loaded into a syringe, and injected in the defect area. The skin incision was closed with wound clips.

### 2.4 Mass spectrometry quantification of pro-inflammatory eicosanoids and SPMs from spinotrapezius and quadriceps muscle tissue

The LC-MS/MS-based lipidomics studies in Figures 1, 2, 6, S2 and S5-S7 were performed as described below: Excised spinotrapezius muscle was homogenized in ice cold PBS using a glass bead homogenizer. Eicosanoids and SPMs were selectively extracted from the homogenized samples by solid phase extraction (SPE) to account for their low concentrations in comparison to lipid species of higher abundance. For this, samples were extracted using an automated C18 SPE manifold (Biotage Extrahera, Uppsala, Sweden). Samples were prepared by depositing homogenized samples on the SPE plate, the sample was rinsed with 800 µL water, followed by 800 µL hexane. The oxylipins were then eluted off from the SPE column with 400 µL methyl formate. The recovered oxylipin fraction was then dried under N_2_ and subsequently reconstituted in methanol to be analyzed by LC/MS. Extracted lipids were resolved using an Agilent Infinity II/6495c LC-MS/MS system. To quantify the SPMs in extract, 10 µL sample was injected onto an Accucore C18 column (100 x 4.6, ThermoFisher, Waltham, MA) and resolved on a 16-minute gradient using water as Solvent A and acetonitrile as Solvent B, both contain 0.1% formic acid. The column was heated to 50°C in a temperature-controlled column chamber and 0.5 mL/min flow rate was used for analysis. Eluted oxylipins were analyzed by Agilent 6495c triple quadrupole mass spectrometer. Instrument parameters (reported below) were optimized using external analytical grade standards and were held consistent over the course of analysis. Oxylipins were analyzed in the negative ion mode using a multiple reaction monitored (MRM) based method. For this, the mass of the target lipid and a characteristic fragment were targeted for detection. Oxylipins were quantified using MassHunter software, where the area under the curve of each identified lipid is calibrated against an external calibration curve. To create the curve, analytical grade standards in the linear range of 0.1 nM-10 nM were created for each standard. The lower limits of detection and upper limits of detection were determined, plotting concentration versus area under the curve. The slope of the linear regression equation was used for calibration of the corresponding analyte. Instrumental parameters were optimized for this lipid class using internal standards and were held constant during the course of the experiment. For each standard, serial dilutions were prepared that cover a broad range of concentrations, typically in the range of 0.01 µM to 1.0 mM, to determine the upper and lower limits of detection (LLOD). It should be noted that the LLOD is at least 10 times the noise level.

**Fig. 1:**
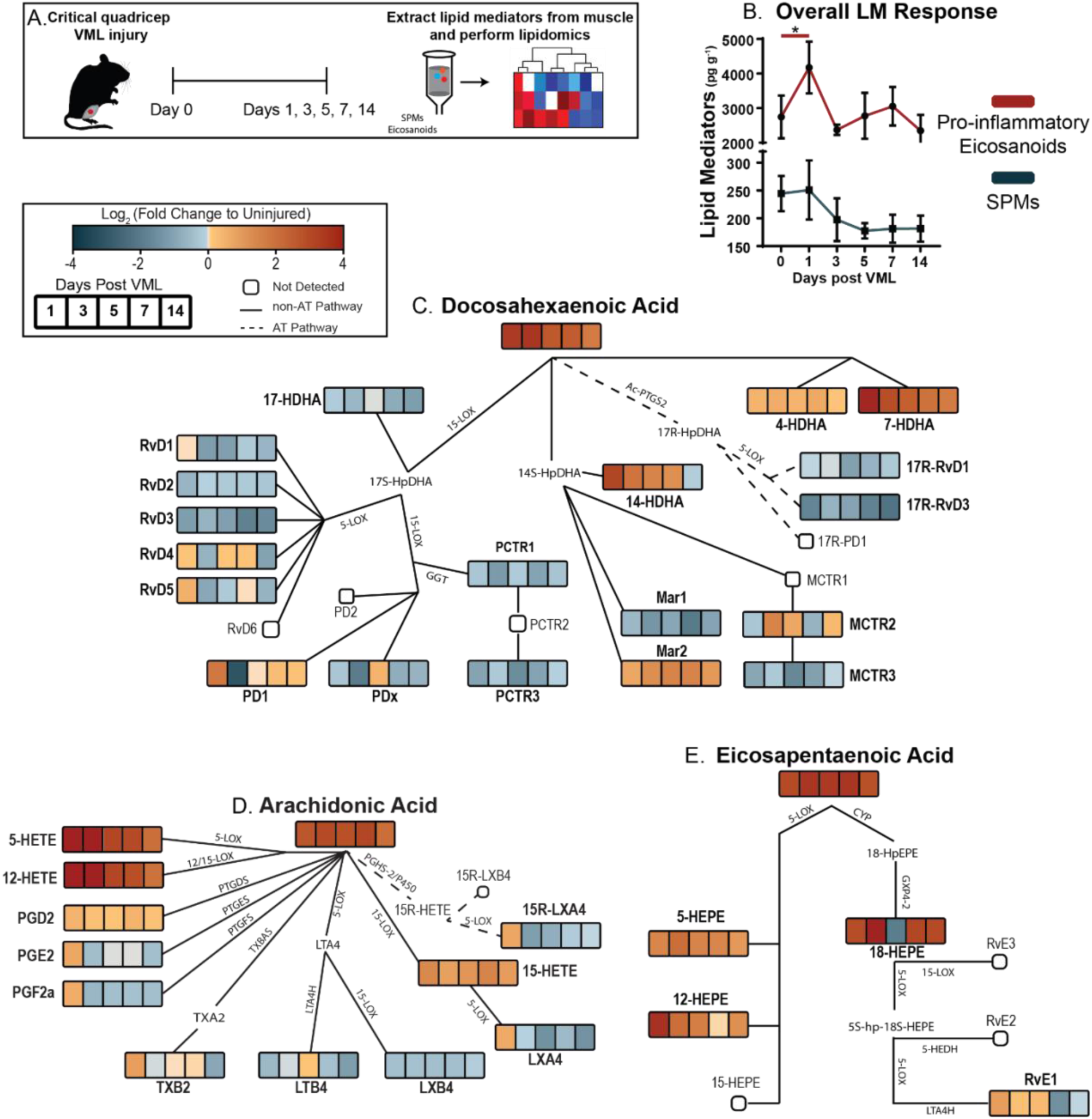
Targeted LC-MS/MS after VML injury reveals impaired pro-inflammatory to pro-resolving lipid mediator switch. A critical VML (3 mm diameter punch) injury was performed to the murine quadriceps muscle and resulting lipid mediators were analyzed via LC-MS/MS at specified timepoints after injury or from uninjured quadriceps (A). (B) Overall lipid mediator response following injury. Lipid mediator concentration heatmaps represented as a fold change between VML and uninjured samples at 1-, 3-, 5-, 7-, and 14-days post-injury are overlaid onto the AA (C), EPA (D), and DHA (E) metabolic pathways. Fold change values range negatively to positively with blue to red color mapping, respectively. Fold change heatmaps created using the average concentration from n=5 animals per condition. Statistical analysis for B was performed using a one-way analysis of variance (ANOVA) with Dunnet’s multiple comparisons; * *p* <0.05; n =5 animals per group.

### 2.5 Mass spectrometry quantification of pro-inflammatory eicosanoids and SPMs from quadriceps muscle tissue

The LC-MS/MS-based lipidomics study in Figure S3 was performed by the following method: Samples of muscle tissue (quadricep) were subjected to solid phase extraction (SPE) followed by a targeted liquid chromatography-tandem mass spectrometry (LC-MS/MS) analysis. Samples were stored at −80°C prior to extraction. Deuterated internal standards including d5-RvD2, d4-LTB_4_, d8-5-HETE, d4-PGE_2_, d5-MaR1, and d5-LXA_4_ (Cayman Chemical) were used to assess extraction recovery and quantification. Mechanically digested samples were centrifuged (3,000 rpm) and supernatants were then subjected to SPE and LC-MS/MS analysis, in part as described previously^17,30^. In short, acidified water (pH 3.5 with HCl) was added to samples immediately prior to SPE using C18 column chromatography. Lipid mediators were eluted in the methyl formate fractions, the solvent was evaporated under a gentle stream of N_2_ gas and the samples were then resuspended in methanol:water (50:50). Samples were then injected using a high-performance liquid chromatograph (HPLC, Shimadzu) equipped with a reverse-phase C18 column (100 mm x 4.6 mm x 2.7 mm; Agilent Technologies) held at 50°C and coupled to a QTrap5500 mass spectrometer (AB Sciex) operating in negative ionization mode and using Analyst software (v1.7). A gradient of methanol-water-acetic acid ranging from 50:50:0.01 (v/v/v) to 98:2:0.01 (v/v/v) was used at a constant flow rate of 0.5 mL/min. Individual lipid mediators were identified using specific multiple reaction monitoring (MRM) transitions and information-dependent acquisition enhanced product ion scanning. Identification was based on matching retention time with authentic standards (Cayman Chemical) run in parallel using specific MRM transitions, as well matching of MS/MS fragmentation ions in selected samples using Sciex OS-Q (v1.7). Quantitation of mediators was then carried out for peaks reaching a signal to noise ratio of least 5, followed by accounting for the extraction recovery of the deuterated internal standards and by interpolation based on calibration curves of external standards for each individual mediator. The data were normalized to muscle tissue weight. The limit of quantification was established for each mediator by determining the lowest amount that could be quantified in replicate injections with a coefficient of variation of less than 20%.

### 2.6 Flow cytometry

To collect tissue for flow cytometry analysis, mice were euthanized via CO_2_ asphyxiation. For analysis of cell composition in spinotrapezius muscles, a 6 mm biopsy punch of muscle tissue centered on the defect was taken, weighed, and digested with 5,500 U/mL collagenase II and 2.5 U/mL Dispase II for 1.5 h in a shaking 37°C water bath. The digested muscles were filtered through a cell strainer to obtain a single cell suspension. Single-cell suspensions were stained for live cells using either Zombie Green or Zombie Violet (BioLegend) dyes in cell-culture grade PBS per manufacturer instructions. Cells were then stained with cell phenotyping antibodies in a 1:1 volume ratio of 3% FBS and Brilliant Stain Buffer (BD Biosciences) according to standard procedures and analyzed on a FACS Aria III flow cytometer (BD Biosciences). Antibodies used for cell phenotyping are listed in Table S1. 30 μL of AccuCheck Counting Beads (Invitrogen) were added per sample for absolute quantification of cell populations. Single, live cells were selected in FlowJo software for subsequent analysis. Myeloid cells were gated as CD11b^+^. Unless indicated otherwise, neutrophils were gated as CD11b^+^ Ly6G^+^ cells. Macrophages were gated as CD11b^+^ CD64^+^ MerTK^+^ cells. Monocytes were gated as CD11b^+^ CD64^+^ MerTK^-^ cells.

### 2.7 High dimensional analysis of flow cytometry data

SPADE is an unsupervised trajectory analysis tool that was designed to map heterogeneous single-cell populations into two dimensions on the basis of similarities across defined markers^31^. SPADE creates a tree structure where nodes represent clusters of cells with similar marker expression. The size and color of each node are relative to the number of cells present and the median marker expression. SPADE was performed in MATLAB, and the source code is available at http://pengqiu.gatech.edu/software/SPADE/. SPADE automatically constructs the tree by performing density-dependent downsampling, cell clustering, linking clusters with a minimum spanning-tree algorithm, and upsampling based on user input. The SPADE tree was generated by exporting uncompensated pre-gated live, single-cell, CD11b^+^CD64^+^MerTK^+^ macrophages, pre-gated live, single-cell, CD11b^+^Ly6G^+^ neutrophils, pre-gated live, single-cell, CD11b^+^CD64^+^MerTK^-^ monocytes, pre-gated live, single-cell, Lin^-^ cells. The markers and/or features used to build the neutrophil SPADE tree were SSC, FSC, CD11b, Ly6G, CD49d, CXCR4, CD62L, CXCR2, CD47, MPO. With this SPADE tree, clusters consistent with the aged neutrophil phenotype were identified. The markers and/or features used to build the macrophage and monocyte SPADE tree were SSC, FSC, CD11b, CD64, MerTK, CD206, Ly6C. With this SPADE tree, clusters consistent with the inflammatory and anti-inflammatory monocyte and macrophage phenotype were identified. The markers and/or features used to build the Lin-SPADE tree were SSC, FSC, Lineage, CD31, CD29, Sca-1, CXCR4. With this SPADE tree, clusters consistent with the MuSC and FAP phenotype were identified. The multiple comparison tool in SPADE was used to identify significantly different populations between treatment groups. The following SPADE parameters were used: apply compensation matrix in FCS header, arcsinh transformation with cofactor of 150, neighborhood size of 5, local density approximation factor of 1.5, maximum allowable cells in pooled downsampled data of 50,000, target density of 20,000 cells remaining, and number of desired clusters of 60.

### 2.8 Fluorescence activated cell sorting (FACS) and *in vitro* cell culture

Fibro-adipogenic progenitor (FAP) cells were isolated from uninjured murine quadriceps by fluorescence activated cell sorting (FACS). Briefly, entire quadriceps muscles were harvested, minced, and digested with 5,500U/mL collagenase II and 2.5 U/mL Dispase II for 1.5 hours on a shaker at 37°C. The digested muscles were filtered through a cell strainer to obtain a single cell suspension. Single cells were then stained with cell phenotyping antibodies (Table S2) in HBSS solution (ThermoFisher) containing 2% FBS. Propidium iodide (PI) was added immediately before sorting for viability. FAPs were identified as PI^-^/CD11b^-^/CD31^-^/Ter119^-^/CD45^-^/Sca-1^+^ and sorted on a FACS Aria III flow cytometer (BD Biosciences). Freshly sorted FAPs were seeded at a density of 10,000 cells/well into 24-well plates (Ibidi) coated with collagen and laminin. FAPs were expanded for 6 days in growth media consisting of DMEM (ThermoFisher), 20% FBS (Atlanta Biologicals), 1% penicillin-streptomycin (PS, ThermoFisher), and 1% GlutaMAX (ThermoFisher). Each day in growth media, cells were supplemented with 2.5 ng/mL b-FGF (Sigma). On day 6, growth media was replaced by differentiation media containing DMEM, 5% horse serum (Atlanta Biologicals), 1% PS, and 1% GlutaMAX. 2.5 ng/mL TGF-β1 (R&D Systems), 100 ng/mL AT-RvD1 (Cayman Chemical), or both were added to FAPs differentiation media to assess the effect of AT-RvD1 on FAPs fibrotic differentiation. 50% media by volume was changed every other day. FAPs were fixed and stained (Table S3) after 4 days of treatment.

### 2.9 Spinotrapezius tissue whole mount immunohistochemistry and confocal imaging

Mice were euthanized 14 days after surgery via CO_2_ asphyxiation. Post-euthanasia, mouse vasculature was perfused with warm saline followed by 4% paraformaldehyde (PFA) until tissues were fixed. The entire spinotrapezius muscle was explanted and permeabilized overnight with 0.2% saponin, then blocked overnight in 10% mouse serum. For immunofluorescence staining, tissues were incubated at 4°C overnight in a solution containing 0.1% saponin, 5% mouse serum, 0.5% bovine serum albumin, and the following conjugated fluorescent antibodies: Alexa Fluor 650 anti-desmin (1:200 dilution, Abcam), Alexa Fluor 650 anti-CD68 (1:200 dilution, Bio-Rad), Alexa Fluor 488 anti-CD206 dye (1:200 dilution, BioLegend). Following immunostaining, tissues were washed four times for 30 min each in 0.2% saponin for the first two washes, 0.1% saponin for the third wash, and PBS for the final wash, and then mounted in 50/50 glycerol/PBS. Mounted samples were imaged on a Zeiss LSM 710 NLO confocal microscope (Objective: 20X / 0.8 NA Plan-Apochromat). Cropped images of 332 × 332 μm at 20X magnification in the injury area were taken for image analysis in ImageJ.

### 2.10 Second harmonic generation (SHG) imaging and analysis

Second harmonic generation (SHG) imaging of whole mount spinotrapezius muscles was performed using a home-built multiphoton microscope, with an optical setup similar to what we reported previously^32^. This system has a Ti:Sapphire femtosecond pulsed laser tuned to 780 nm, with the power of the beam adjusted by a half wave retarder and polarizing beam splitter, then rapidly modulated by a Pockels cell attenuator and scanned over the specimen. The back-detected emission light is sent to a photon multiplier tube with a bandpass filter of 390/18 to capture SHG images of whole mount spinotrapezius muscle. SHG images (512 x 512 pixels) of spinotrapezius muscle structure were collected with linearly polarized excitation using a half-waveplate sequentially rotated to 0°, 90°, and 180° to enhance contrast for evaluating muscle fiber orientation. For all samples, the edges of each volumetric muscle loss defect area were imaged to create 3 polarized images from similar regions of the defect areas. We used a polarization state analyzer to measure the Stokes parameters for each excitation angle and correct for polarization distortion in the focus. We analyzed SHG images in the Fourier space, which represents the periodicity and angle of structures in the samples. We then used an angular Fourier filter (AFF) developed previously^33^ to analyze the image content across different orientation angles, where image content that is spread across multiple angles suggests less organized tissue. Because muscle fibers have very directionally organized structures in their healthy state, this metric can assess the state of the tissue during regenerative processes.

### 2.11 Quadricep tissue histology, immunostaining, and confocal imaging

Tissue histology and immunostaining were performed as previously reported^28^. Briefly, cryosections (CryoStar NX70 Cryostat) were taken at 10 μm thickness and stained with hematoxylin and eosin (H&E) (VWR, 95057-844, -848) according to the manufacturers’ instructions. Before antibody staining, tissue sections were blocked and permeabilized using blocking buffer (5% BSA, 0.5% goat serum, 0.5% Triton-X in PBS) for 30 min and an additional wash with Goat F(ab) anti-mouse IgG (Abcam; ab6668, 2 μg/mL in blocking buffer) was performed for 1 h. Samples were washed between steps with 0.1% Triton in PBS. Primary antibodies were diluted in blocking buffer at 1:200 for dystrophin (Abcam; ab15277) and von Willebrand factor (vWF) (Abcam; ab6994). Primary antibody for embryonic myosin heavy chain (eMHC) (DSHB, F1.652) was diluted 1:10 in blocking buffer. All primary antibodies were incubated for 1 h. Secondary antibodies were conjugated to Alexa Fluor 488 (ThermoFisher; Ms: A-11029, Rb: A-11034), 555 (ThermoFisher; Ms: A-21424, Rb: A-21429), or 647 (ThermoFisher; Ms: A-21236, Rb: A-21245). All secondary antibodies were diluted 1:250 in blocking buffer and incubated for 30 min. Slides were mounted with Fluoroshield Mounting Medium with DAPI (Abcam; ab104139) and stored at 4°C. Immunofluorescence images were taken on Nikon W1 Spinning Disk Confocal microscopes at 20X (0.75 numerical aperture, working distance of 1.0 mm) and stitched together with Nikon Elements AR.

### 2.12 Pain Behavior Testing

*Dynamic Weight Bearing (DWB) –* DWB assesses spontaneous nociception in rodents^34,35^. The DWB device (Bioseb, Vitrolles, France) consists of a Plexiglass chamber with a pressure sensor matrix in the floor and a top-view camera mounted in the ceiling. The pressure sensor was tared and calibrated weekly during the study period. Mice were randomly assigned to an acquisition order which was kept constant at each timepoint. For data acquisition, mice were placed alone in the acquisition chamber for 5 minutes each, without any habituation period before recording. Feces and urine were wiped from the chamber with a dry Kimwipe between each mouse. Data were processed in the Bioseb ADWB2 software (version 2.2.6). Scoring was performed on immobility postures (immobility threshold: 700 ms). Week 4 DWB measurements were not acquired due to sensor malfunction.

*Von Frey (VF) –* Mechanical allodynia, or hypersensitivity to a benign mechanical stimulus, is a hallmark of neuropathic pain^36–38^. It is commonly assessed in rodents using von Frey filaments or hairs which are calibrated to exert a specific force onto the subject’s skin^39^. Researchers were blinded to experimental groups for the duration of the study. The von Frey setup consists of a 2x4 grid of polycarbonate boxes set on an elevated platform with a wire mesh floor to allow for full access to the paws. Mice were randomly assigned to grid locations, and they were placed in the same location for each timepoint. Before the study and at each timepoint, mice were acclimated to the setup for approximately 10 minutes until grooming and exploration behaviors were minimal. The mid-plantar region of each hind-paw was then probed with a series of von Frey filaments (Bioseb, Vitrolles, France), exerting a force of 0.07, 0.16, 0.4, 0.6, 1.0, or 1.4 grams to the paw, in ascending order. The series was repeated for a total of two rounds. Positive withdrawals were recorded as a 1, and no response was recorded as a 0. The average withdrawal response for each mouse’s left and right limbs were calculated by averaging together the recorded responses from all six filaments.

### 2.13 *In vivo* isometric torque analysis

Quadriceps function was assessed by measurement of isometric torque production about the knee, according to an adapted protocol from previous studies^40^. Briefly, mice were anesthetized with 2% isoflurane. Hindlimbs were shaved and a 1 cm incision was made in the skin on each side to expose the quadriceps and medial vasculature and motor neurons. The femoral nerve was isolated from the surrounding connective tissue. Mice were fixed in place in a supine position on the testing box and the ankle of one limb at a time was secured to a force transducer. The femoral nerve was stimulated with a nerve cuff attached to a Grass S11 Stimulator set to 0.2 ms pulse duration, 5.7 ms pulse interval, and 500 ms train duration. The stimulus was directed through a Grass Stimulus Isolation Unit (SIU) before connecting to the nerve cuff. Force data was recorded using a data acquisition board (USB-1608G) and LabView software. Voltage was varied until maximal torque was measured, then 3 trials were recorded. Data analysis was done using MATLAB (Mathworks, Natick, MA). The find peaks function was used to pick out the 3 peaks from the testing file. Voltage values were converted to force and then multiplied by an average mouse tibia length (2 cm) to get torque about the knee.

### 2.14 Statistical analysis

Data is presented as means ± SEM unless otherwise noted. Statistical tests were conducted using GraphPad Prism 8. Two-tailed unpaired *t*-tests were performed for two-group comparisons. For comparison of more than two groups, one-way ANOVA with Tukey’s *post hoc test* used for multiple comparisons. For grouped analyses, two-way ANOVA with Sidak’s *post hoc* test was used for multiple comparisons. Partial Least Squares Discriminant Analysis (PLSDA) of quadriceps lipidomics data was performed with Python 3 using the scikit-learn module^41^. Separate PLSDA models were generated for each timepoint to better assess differential effects of treatments on lipid mediator dynamics. Briefly, lipid mediator concentrations were normalized to muscle weight and then standardized by the mean and standard deviation across treatment groups. Significance levels for all analyses were set at * *p* <0.05, ** *p* <0.01, *** *p* <0.001, **** *p* <0.0001. Investigators were blinded to the experimental groups when analyzing the data. Differences in sample sizes between groups are due to instrument errors causing no detection in cells or metabolites; however, these studies remain adequately powered for statistical analysis.

## 3. RESULTS

### 3.1 Lipid mediator class-switching is dysregulated after volumetric muscle loss (VML) injury

The lipid mediator response to sub-critical, self-healing muscle injuries has been well characterized^17–19^. These studies show that after acute muscle injury an increase in pro-inflammatory lipid mediators quickly returns to baseline levels. This decrease in pro-inflammatory lipid levels occurs in parallel to an increase in the concentration of specialized pro-resolving lipid mediators in the acute injury environment^17^. We performed liquid chromatography-mass spectrometry (LC-MS) to assess pro-inflammatory eicosanoids and SPMs in an established murine quadriceps animal model in which non-healing, critical-sized (VML) injury is achieved (Fig. 1A)^28,42^. This LC-MS-based analysis showed that the lipid mediator response of a non-healing injury deviated from that of an acute injury. In VML, an initial and significant spike in the concentration of pro-inflammatory lipid mediators returned to baseline levels by day 3 post-VML injury (Fig. 1B). Although the pro-inflammatory lipid mediator concentration was not sustained after VML injury, the concentration of pro-resolving lipid mediators did not increase after injury. Rather, SPM concentration showed a steady decrease up to 14 days following VML injury (Fig. 1B).

Visualization of changes to the omega-3 fatty acid metabolic networks after VML are displayed using a heatmap; the lipid mediator concentration data for each detected analyte is represented as a fold change normalized to uninjured quadriceps muscle and overlayed onto the corresponding metabolite (Fig. 1C-E). Our analysis of the arachidonic acid (AA) metabolome revealed a short-lived increase in production of pro-inflammatory lipid mediators 1 day after VML injury, which returned to baseline levels at 3 days post-VML, except for PGD2, which remained elevated throughout the 14-day timeline. There was also a delayed production spike of LTB4 at 5 days post-VML (Fig. 1D). In the pro-resolving lipid mediator compartment of the AA metabolome, LXA4 showed a slight increase in concentration at day 1 post-VML that then decreased to levels below that of uninjured quadriceps at days 3 through 14 post-injury while LXB4 levels were decreased at all timepoints (Fig. 1D). Most pro-resolving lipid mediators are derived from eicosapentaenoic acid (EPA) and docosahexaenoic acid (DHA). Analysis of these networks showed an inability of the local VML niche to produce pro-resolving lipid mediators in a sustained manner (Fig. 1C and E). The concentration of EPA-derived RvE1 was elevated at days 1, 3 and 5 post-VML before dropping to levels below that of uninjured muscle at day 7 post-VML (Fig. 1E). The concentration of DHA-derived Mar2 and PD1 both showed a sustained elevation out to 14 days post-VML (Fig. 1C). However, apart from those few exceptions, the analysis of the pro-resolving lipid mediator concentrations demonstrated a decrease in lipid mediator concentration to levels below that of uninjured muscle at days 3 through 14 post-VML (Fig. 1B).

To characterize the activation or downregulation of major enzymatic steps in the production of downstream pro-inflammatory and pro-resolving lipid mediators, we analyzed the monohydroxy pathway intermediate concentrations. An analysis of these intermediates in the AA metabolome revealed that the conversion of AA to 15-HETE via 15-LOX was upregulated after VML as indicated by a significant increase in both the concentration of AA and 15-HETE (Fig. S2A). However, the concentration of pro-resolving LXA4 showed a downward trend after VML injury indicating that the ability of the injury milieu to convert 15-HETE to LXA4 was impaired (Fig. S2A). Additionally, the significant increase in the concentration of PGE2 and TXB2 at 1-day post-VML before returning to baseline levels showed a transient activation of COX acting on AA (Fig. S2A). The main substrates for pro-resolving lipid mediators, DHA and EPA, showed a sharp and sustained increase in concentration with DHA concentration being significantly increased at days 1 and 3 post-VML and EPA concentration being significantly increased at days 3, 5 and 7 post-VML (Fig. S2B and C). Despite the initial increase in DHA, a downward trend of the concentration of 17-HDHA showed that the conversion of DHA by 15-LOX to 17-HDHA does not occur after VML injury (Fig. S2B). This was further supported by the concentrations of the downstream lipid mediators in this pathway with the concentration of RvD3 being significantly decreased and the concentrations of RvD1 and PD1 remaining unchanged (Fig. S2B). Additionally, 14-HDHA, the pathway marker for 12-LOX acting on DHA, showed a significant increase in concentration at day 1 post-VML; however, the downstream lipid MAR1 exhibited a steady decrease in concentration with significance at 7 days post-VML (Fig. S2B). The conversion of DHA to the 17R-series of Resolvins was also downregulated as shown by the significant decrease in the concentration of 17R-RvD3 at days 1, 7 and 14 post-VML (Fig. S2B). Similarly to DHA, even with a more sustained increase in the concentration of EPA, the conversion of EPA to bioactive lipid mediators was dysregulated as evidenced by the unchanged concentration of RvE1 (Fig. S2C). These results are further supported by comparing the lipid mediator concentrations between a sub-critical injury to a critically sized VML injury (Fig. S3), where there was an increase in AA-derived pro-inflammatory Prostaglandin concentrations (Fig. S3B) and an accumulation of monohydroxy intermediates (e.g. 17-HDHA, 14-HDHA) in SPM pathways rather than their downstream SPM products in critical VML compared to sub-critical (Fig. S3C and D).

Taken together, these data suggest that the normal lipid mediator class switch from pro-inflammatory to pro-resolving lipid mediators that occurs during self-limited inflammation and routine muscle healing^19^, is dysregulated after a VML injury. These observations led us to design a pro-resolving therapeutic strategy to increase the production of SPMs after VML as a functional regenerative therapy.

### 3.2 Design of a PEG-4MAL hydrogel to promote pro-resolving lipid mediator biosynthesis

To engineer an injectable delivery platform that activates local pathways of inflammation resolution, we used a synthetic hydrogel platform based on a four-arm PEG macromer functionalized with maleimide groups for local, controlled delivery of SPMs^26,27^. In this synthetic design, the maleimide in PEG-4MAL macromers were functionalized with thiol-containing cell-adhesive peptide (RGD) via Michael-type addition (Fig. 2A). The functionalized macromer was then crosslinked through the reaction of the remaining maleimides with cysteine-flanked protease-degradable peptides to yield a crosslinked hydrogel that is enzymatically degraded *in vivo*^27,43^. This crosslinking reaction is performed in the presence of 100 ng AT-RvD1 (17R-Resolvin D1), a dosage chosen based on previous studies of repeat systemic Resolvin delivery^19^, to encapsulate this SPM. This process generates a hydrogel that will locally release the pro-resolving molecule and activate the resolution of inflammation (Fig. 2A). This synthetic hydrogel construct was characterized in previous studies, and bioactive AT-RvD1 was found to be released within the first 24 hours making it bioavailable to the infiltrating pro-inflammatory cells early in the inflammatory cascade^26^.

**Fig. 2:**
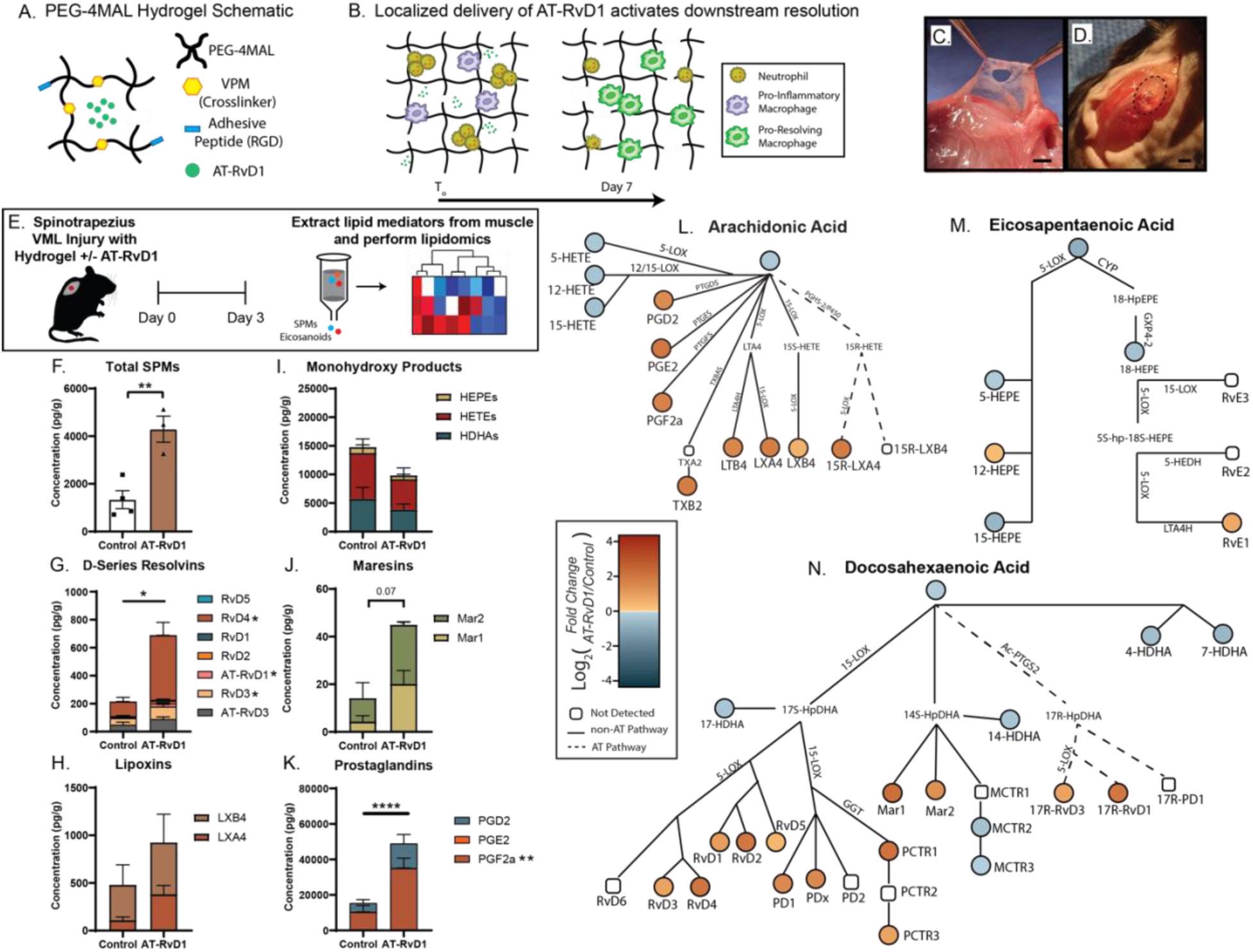
Local delivery of AT-RvD1 via PEG-4MAL hydrogel increases the concentration of specialized pro-resolving mediators. (A) Protease degradable hydrogels were designed to locally deliver AT-RvD1 by functionalizing PEG-4MAL with an adhesive peptide (RGD) and crosslinking it with an MMP-cleavable crosslinker (VPM) in the presence of AT-RvD1. (B) The downstream inflammatory cascade is reprogrammed towards resolution via the early and local presentation of AT-RvD1. (C and D) A volumetric muscle loss injury is induced in the spinotrapezius muscle, and the hydrogel injected to fill the injury void. (E) Modulation of the pro-inflammatory Eicosanoid and Specialized pro-resolving lipid mediator landscape was analyzed via LC/MS-MS three days post-VML. (F-K) Quantification of lipid mediator family concentrations between AT-RvD1 and control hydrogels. (L-N) The fold change difference 3 days post-VML between AT-RvD1 hydrogel treatment and control hydrogel was calculated and converted to a heatmap. This heatmap is overlayed onto the Arachidonic Acid (L), Eicosapentaenoic Acid (M) and Docosahexaenoic Acid (N) metabolic pathways. Statistical analyses were performed using two-tailed *t* tests; Significance values are for the comparison between total groups (e.g. D-Series Resolvins and Prostaglandins) not individual sub-species. Asterisks next to individual lipid species indicate significance values between the control and AT-RvD1 groups for that species. * *p* < 0.05, ** *p* < 0.01, **** *p* <0.0001; n = 3 to 4 animals per group.

A full-thickness VML defect was in the mouse spinotrapezius muscle was used to test the efficacy of early hydrogel-based presentation of AT-RvD1 to resolve chronic inflammation (Fig. 2B). This model has been used in previous studies to characterize the inflammatory and muscle healing response to various therapeutics^44,45^. When subjected to a 2 mm full-thickness defect (Fig. S4A), the spinotrapezius muscle showed the hallmarks of a VML injury, including the inability of the wound to heal (Fig. S4B-D), a severely disrupted vascular structure (Fig. S4E-H), and a persistent inflammatory response (Fig. S4I-L). The hydrogel was injected for *in situ* gelation directly into the VML defect (Fig. 2D). This model allows us to investigate the effects of the therapeutic platform on the cellular environment and lipid-mediator biosynthesis while also allowing for whole-mount microscopy to observe muscle fiber regeneration across the entire thickness of the muscle, an advantage of the model compared to the larger pre-clinical VML model in the quadriceps^29^.

### 3.3 AT-RvD1-loaded hydrogel delivery has broad pro-resolving metabolic effects

We compared the ability of local hydrogel-mediated AT-RvD1 delivery to a control hydrogel (without AT-RvD1) to influence downstream production of key SPMs in muscle tissue using mass spectrometry-based lipidomics at 3 days post-VML injury (Fig. 2E). With this method, we detected significant increases in the concentration of total SPMs including AT-RvD1 and multiple other D-series Resolvins in the AT-RvD1 hydrogel-treated group compared to the control hydrogel-treated group (Fig. 2F and G). Additionally, hydrogel delivery of AT-RvD1 shifted lipid metabolism towards production of downstream bioactive SPMs with a decreased trend in the concentration of monohydroxy intermediates and increased trend in Maresins and Lipoxins (Fig. 2F-J). Interestingly, we also observed a significant increase in Prostaglandin concentration which has been identified as beneficial to efficacious muscle fiber regeneration^46^ (Fig. 2K). These differences, as compared to control hydrogel, were seen at 3 days post-VML in the AT-RvD1 loaded hydrogel treated group. These data indicate that local delivery of a single AT-RvD1 dose can exert significant pro-resolving effects at a stage when the environment is no longer being supplied with external AT-RvD1.

To visualize the effects of the pro-resolving hydrogel on the greater metabolic network from which pro-inflammatory eicosanoids and SPMs are derived, a fold change (AT-RvD1 versus control) heatmap was overlayed onto the DHA, EPA and AA metabolic networks (Fig. 2L-N). This visualization showed that many metabolic pathways were activated with AT-RvD1 delivery as compared to a control hydrogel. In the DHA pathway, the most upregulated lipid mediators, Resolvin D4 (RvD4), Mar1, and AT-RvD1, are from different pathways within the metabolic network (Fig. 2N). This shows that the delivery of AT-RvD1 can increase the *in vivo* biosynthesis of a wide range of SPMs across the metabolic network.

### 3.4 AT-RvD1 delivery decreases the persistent “aged” neutrophil subpopulation

To examine the impact of modulating SPM metabolism in the local VML injury niche on the immune response, we performed flow cytometry and unsupervised trajectory analysis of VML injuries in the spinotrapezius muscle at days 1, 3, and 7 using the Spanning Tree Progression of Density Normalized Events (SPADE) algorithm (Fig. 3A). The initial neutrophil cell compartment is essential in priming the inflammatory response towards either resolution or persistence^47,48^. In the case of persistent inflammation produced by critical VML, neutrophils can exhibit an aged phenotype prone to develop neutrophil extracellular traps (NET) that promote further inflammation and activate fibrotic pathways^49^. We have previously used SPADE to identify these rare neutrophil subsets^47,50^.

**Fig. 3:**
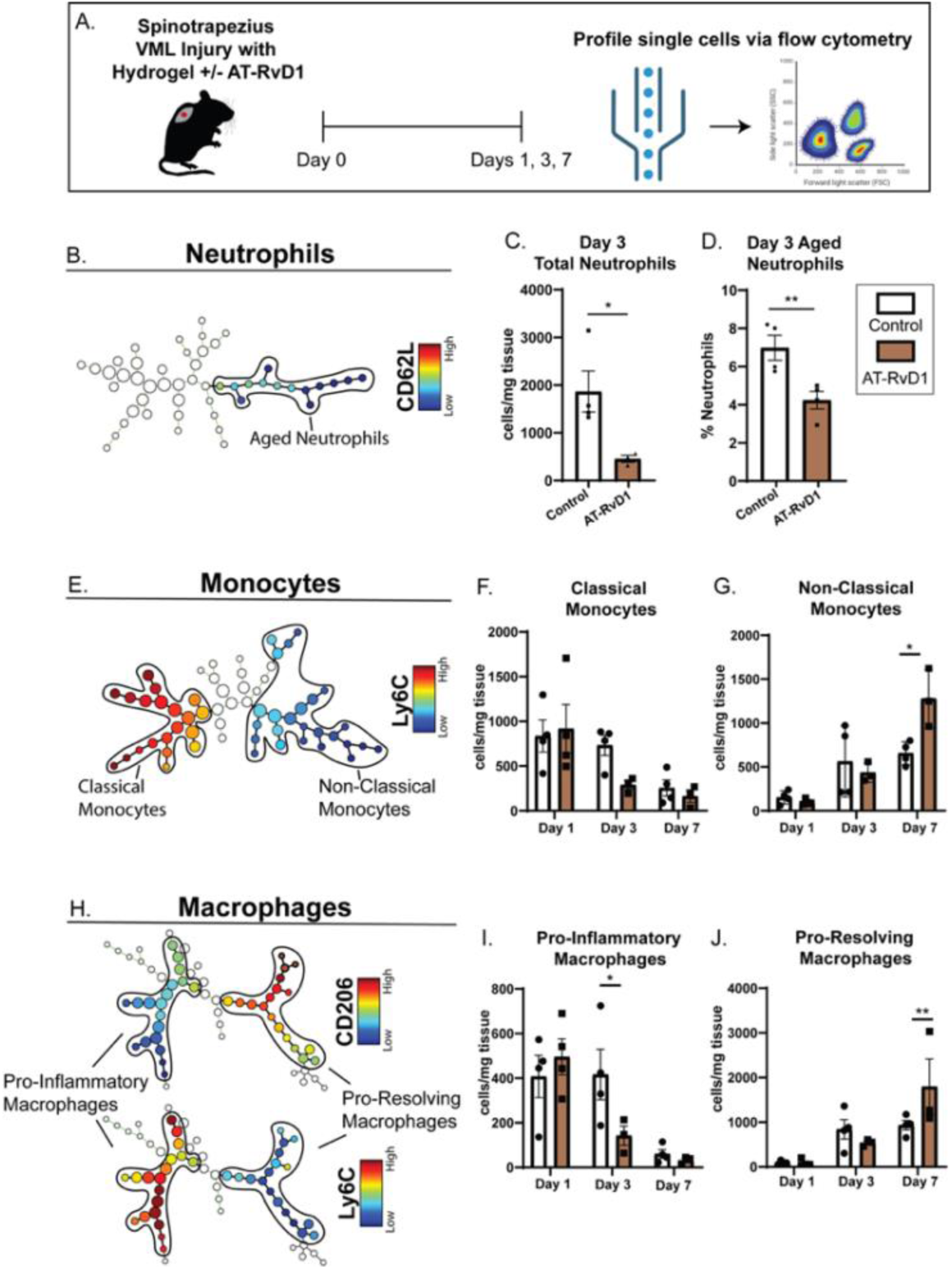
AT-RvD1 loaded hydrogels delivered to a VML injury limits neutrophil accumulation and induces a downstream pro-regenerative cellular niche. (A) Changes in immune responses after local delivery of control or AT-RvD1 loaded hydrogels were assessed by performing flow cytometry at days 1, 3, and 7 after spinotrapezius VML injuries. (B) Traditionally gated CD11b^+^Ly6G^+^ neutrophils were analyzed via SPADE to identify an aged neutrophil trajectory. (C and D) AT-RvD1 was able to reduce the total concentration of neutrophils in the tissue (C) and was also able to limit the polarization of neutrophils to the aged phenotype (D). (E) CD11b^+^CD64^+^Mertk^-^ monocytes were analyzed via SPADE to identify classical and non-classical monocyte phenotypes, based on high or low Ly6C expression, respectively. (F and G) Quantification of classical and non-classical monocyte subsets in the muscle tissue across timepoints between AT-RvD1-loaded and control hydrogel groups. (H) CD11b^+^CD64^+^Mertk^+^ macrophages were analyzed via SPADE to identify Ly6C^Hi^CD206^Low^ pro-inflammatory M1 and Ly6C^Low^CD206^Hi^ pro-resolving M2 macrophages. (I and J) Quantification of pro-inflammatory and pro-resolving macrophage subsets in muscle tissue across timepoints between AT-RvD1-loaded and control hydrogels. Statistical analyses were performed using two-tailed t tests (C-D) or a two-way analysis of variance (ANOVA) with Sidak’s multiple comparisons and the Geisser-Greenhouse correction to account for the differences in variability amongst groups (F-G, I-J); * *p* < 0.05, ** *p* < 0.01; n = 3 to 4 animals per group.

Visualization of the CD11b^+^Ly6G^+^ tree revealed a cluster of neutrophils consistent with the “aged” phenotype (Fig. 3B). This aged phenotype was characterized by low expression of CD62L and high expression of CD49D and CD184. Additionally, this subset also has a high expression of CD47, a marker that allows cells to evade phagocytosis, and myeloperoxidase (MPO), a key component of NETs. With local hydrogel delivery of AT-RvD1, the total concentration of neutrophils in the muscle tissue at 3 days post-VML was significantly decreased compared to the control hydrogel (Fig. 3C). Furthermore, the pro-resolving hydrogel reduced the polarization of activated neutrophils toward a pro-inflammatory aged phenotype (Fig. 3D). This modulation of neutrophil infiltration and phenotype indicates that local AT-RvD1 promotes cellular resolution of inflammation.

### 3.5 Local AT-RvD1 delivery promotes pro-regenerative polarization of mononuclear phagocytes

The timely resolution of inflammation is driven by the shift from pro-inflammatory (M1) macrophages to pro-resolving or pro-regenerative (M2) macrophages^51^. This shift involves the recruitment of either inflammatory or “classical” monocytes that differentiate into M1 macrophages or recruitment of anti-inflammatory or “non-classical” monocytes that differentiate into M2 macrophages. Additionally, M1 macrophages can polarize to M2 macrophages that produce large amounts of SPMs and further promote inflammation resolution^51^.

The effect of local AT-RvD1 delivery on the pro-resolving shift in immune cells was quantified in the spinotrapezius muscle after VML injury via SPADE analysis. A monocyte SPADE tree was constructed using CD11b^+^CD64^+^Mertk^-^ monocytes and clustering identified a separation between Ly6C^high^ inflammatory and Ly6C^low^ anti-inflammatory monocytes (Fig. 3E). Local delivery of AT-RvD1 significantly increased the concentration of anti-inflammatory monocytes at day 7 post-VML (Fig. 3F and G). Similarly, a macrophage SPADE tree of CD11b^+^CD64^+^Mertk^+^ macrophages (Fig. 3H) revealed a shift in the proportion of CD206^-^Ly6C^high^ M1 and CD206^+^Ly6C^low^ M2 macrophages. Local AT-RvD1 delivery promoted an M1 to M2 macrophage switch resulting in a significant decrease in M1 macrophages at day 3 and a significant increase in M2 macrophages at day 7 post-VML (Fig. 3I and J). Taken together, these data indicate that local AT-RvD1 presentation shifts the distribution of mononuclear phagocytes towards a pro-regenerative phenotype.

### 3.6 Pro-resolving injury micro-environment increases the skeletal muscle regenerative capacity

To analyze the effect of local AT-RvD1 hydrogel delivery on the local pool of the two main progenitor cell populations in muscle – muscle satellite/stem cells (MuSCs) and fibro-adipogenic progenitor (FAP) cells, we quantified the concentration of these cells at days 1, 3 and 7 post-VML in the spinotrapezius muscle (Fig. 4A and B). A SPADE tree of all lineage-negative cells was constructed, and clustering identified distinct trajectories comprised of FAPs (CD31^-^SCA1^+^) or MuSCs (CD31^-^SCA1^-^CD29^+^CD184^+^). FAPs and MuSCs work in concert to provide the necessary extracellular matrix (ECM) structure and muscle fiber formation, respectively, for optimal regeneration to take place^52^. When quantifying the SPADE-identified population of MuSCs and FAPs, AT-RvD1 hydrogel treatment significantly increased both populations compared to control hydrogel at 7 days post-VML (Fig. 4A and B).

**Fig. 4:**
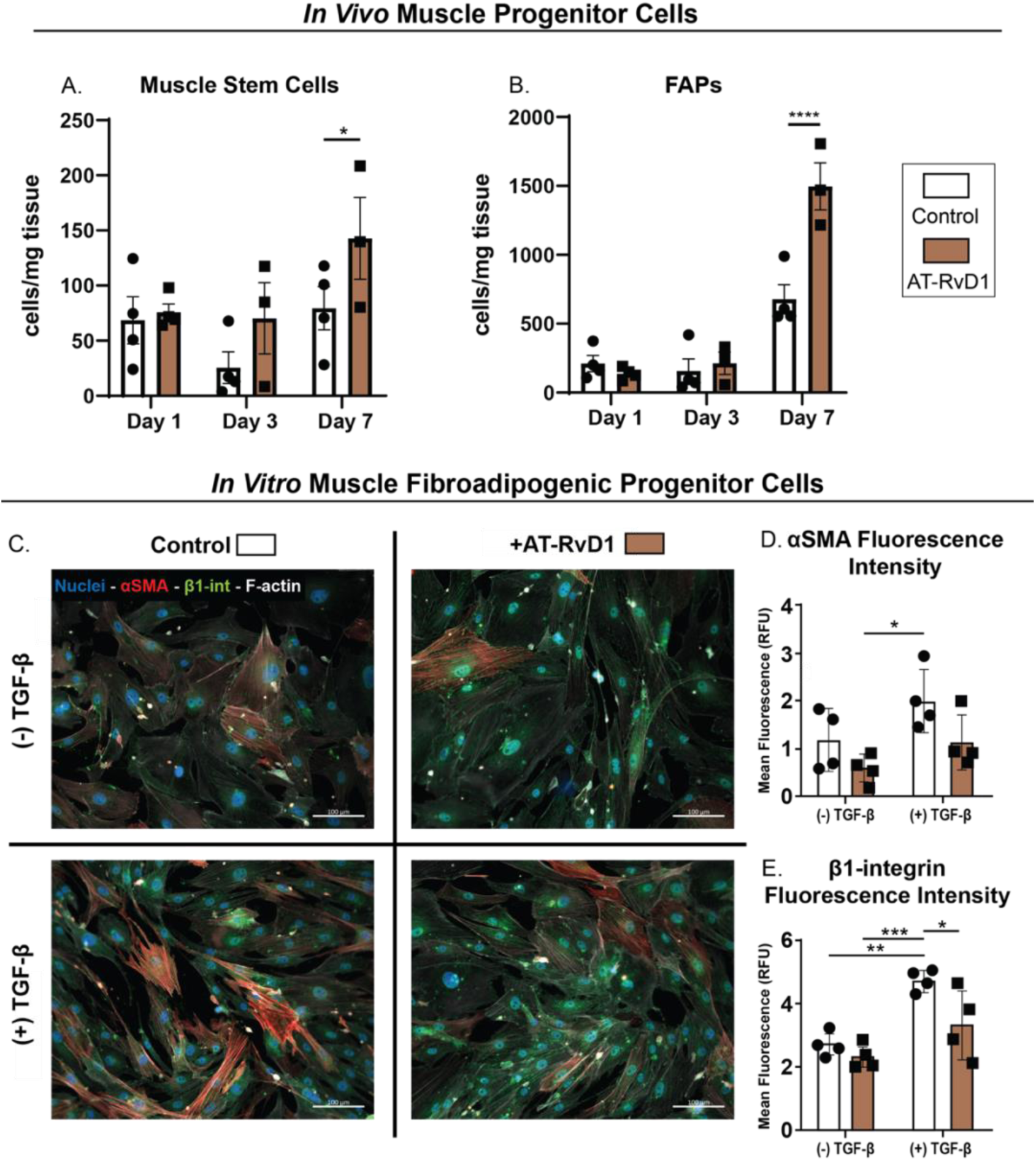
AT-RvD1 acts on muscle progenitor cells to increase the skeletal muscle regenerative capacity post-VML. (A and B) Traditionally gated Lineage^-^ cells were analyzed via SPADE to identify MuSCs (Lin^-^CD31^-^SCA1^-^CD29^+^CXCR4^+^) and FAPs (Lin^-^CD31^-^SCA1^+^). Their concentrations were quantified in the muscle tissue at 1-, 3-, and 7-days post-injury. (C) FACS-sorted FAPs from muscle tissue were stained with aSMA and β1-int to track their fibrotic differentiation in the presence of AT-RvD1 with or without TGF-β stimulation. (D and E) Fluorescence intensity of aSMA (D) and β1-int (E) show that AT-RvD1 directly interacts with FAPs to decrease their fibrotic differentiation. Statistical analyses were performed using a two-way analysis of variance (ANOVA) with Sidak’s multiple comparisons and the Geisser-Greenhouse correction to account for the differences in variability amongst groups; * *p* < 0.05, ** *p* < 0.01, *** *p* <0.001 and **** *p* <0.0001; n = 3 to 4 animals per group.

Previous studies have shown that Resolvins can have a direct effect on MuSCs to promote proliferation, differentiation, and myotube formation^18,19^. However, the effects that SPMs have on FAPs have yet to be explored. Thus, we isolated FAPs from naïve muscle tissue using fluorescence activated cell sorting (FACS) and cultured them for downstream analysis (Fig. 4C-E). Once the FAPs were expanded in culture, we compared the fibrotic differentiation of the cells treated with or without AT-RvD1 in combination with or without TGF-β (a growth factor that drives FAPs towards a fibrotic phenotype). FAPs were stained with Hoechst (blue) and phalloidin (white) to visualize the cells along with α-SMA (red) and β1-integrin (green) to quantify fibrotic differentiation (Fig. 4C). TGF-β stimulation increased the fluorescent intensity of both α-SMA and β1-integrin. Treatment of the TGF-β stimulated FAPs with AT-RvD1 attenuated α-SMA expression to levels equivalent to unstimulated FAPs (Fig. 4D). Additionally, the expression of β1-integirn by TGF-β-stimulated FAPs was significantly decreased when treated with AT-RvD1 (Fig. 4E). This ability of AT-RvD1 to act directly on FAPs and limit expression of both α-SMA and β1-integrin demonstrates the anti-fibrotic actions of AT-RvD1 treatment. Taken together, our results indicate that in addition to acting on immune cells to promote resolution, AT-RvD1 can promote MuSC proliferation and differentiation as well as directly limit FAP pro-fibrotic differentiation.

### 3.7 AT-RvD1 hydrogel delivery increases muscle regeneration and recovers collagen organization

The spinotrapezius pre-clinical VML model can be whole mounted for full thickness confocal microscopy. This allows us to analyze the extent of VML defect closure without treatment and with a control (empty) and AT-RvD1-loaded hydrogel treatment (Fig. 5A). At 14 days post-VML injury, we stained the spinotrapezius muscle for desmin (red; muscle fibers), CD68 (blue; macrophages) and CD206 (green, M2 surface marker) (Fig. 5B-D). Untreated defects remained significantly large (> 3 mm^2^) and devoid of repair tissue. In contrast, defects treated with either control hydrogel or AT-RvD1 hydrogel exhibited a significantly smaller defect area with aligned desmin^+^ fibers at the periphery of the defect (Fig. 5E). Strikingly, defects treated with AT-RvD1 loaded hydrogels exhibited a defect area with near complete closure that was 7 times smaller than those treated with control hydrogels and more than 30 times smaller than untreated defects (Fig. 5E). CD68 and CD206 staining also showed that M2 macrophages (showing expression of both surface markers) tended to cluster around newly generated fibers. We observed that these cells are constrained to the outer edges of the defect without much infiltration into the center of the defect with a control hydrogel (Fig. 5C). Notably, with AT-RvD1 loaded hydrogels, M2 macrophages spread throughout the defect area (Fig. 5D). Additionally, SHG imaging was leveraged to produce data within one mean-free-path from the surface of untreated, control hydrogel, AT-RvD1 hydrogel and healthy (intact) mouse spinotrapezius whole mount slides. Angular Fourier filter (AFF) was used for the polarimetric analysis of muscle structures^33^. Representative polarization-encoded presentation of the data at 14 days post spinotrapezius VML of the experimental and control groups is shown (Fig. 5F-M). The AT-RvD1 treated VML injury displayed more developed and well-defined collagen and muscle fiber structures, similar to that of the healthy muscle (Fig. 5F-M). AFF is able to detect the presence of a loose and less-organized collagen network versus one that is more defined and mature^33^. A heatmap of the AFF can be used to represent the dominant peaks of an organized collagen and muscle fiber network (Fig. 5N-Q). In both untreated and control hydrogel samples, the AFF analysis did not produce any significant peak but rather showed angularly scattered peaks (Fig. 5N and O). In contrast, healthy and AT-RvD1 hydrogel-treated VML muscle showed strong peaks that followed a sinusoidal trend with the input polarization (Fig. 5P and Q). These results indicate that, for injury without treatment or treated with a control hydrogel, collagen network disruption persists at 14 days post-VML injury, with little muscle fiber regeneration. Strikingly, when treated with AT-RvD1 hydrogels, the regenerated muscle recovered its organization at 14 days post-VML injury and had well-defined collagen fibers similar to that of healthy muscle along with multiple muscle fibers in the defect region. Additionally, as a measure of muscle fiber compactness, percent area of muscle fiber was quantified. Muscle fiber area for AT-RvD1 hydrogel-treated muscle was equivalent to healthy muscle and significantly higher than both control hydrogel-treated and untreated VML muscle (Fig. 5R). Thus, AT-RvD1 hydrogel-treated muscle, akin to healthy muscle, showed compact muscle fiber organization. Taken together, these data show that AT-RvD1-loaded hydrogel treatment after VML injury results in significant muscle defect closure characterized by compact muscle fibers with a defined and organized collagen and muscle network.

**Fig. 5:**
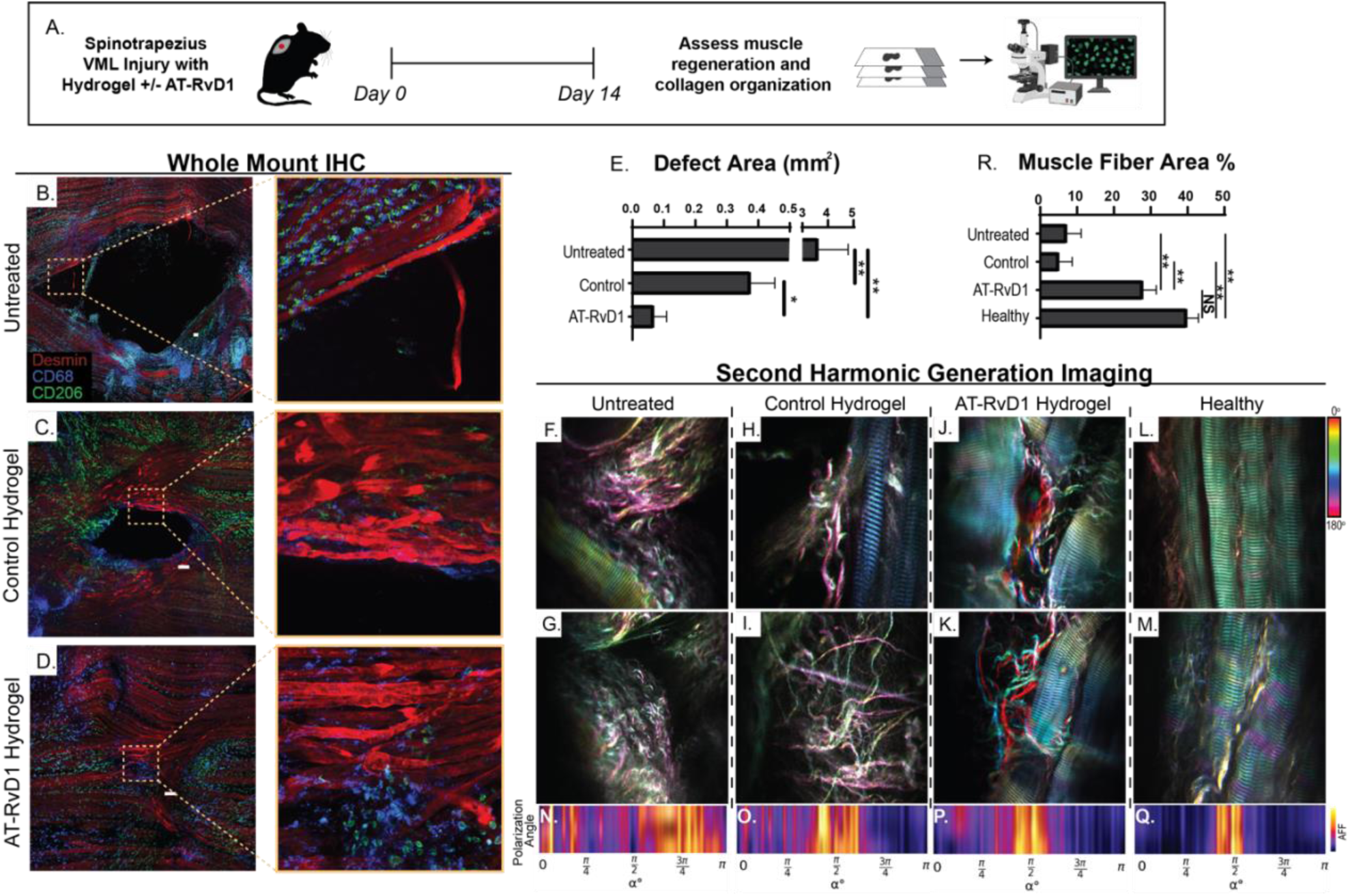
AT-RvD1 loaded hydrogel treatment improves muscle regeneration and recovers collagen organization 14 days post-VML injury. (A) Mice received a spinotrapezius VML injury that was treated with either a control or AT-RvD1-loaded hydrogel. Mice were taken down at day 14 to assess muscle regeneration and collagen organization via whole-mount IHC and SHG imaging, respectively. (B-D) Representative images of whole-mount spinotrapezius confocal miscroscopy 14 days post-VML injury. Desmin (red) stains muscle fibers, CD68 (blue) stains macrophages, CD206 (Green) identifies CD68^+^ M2 macrophages. (E) Quantification of the defect area (area not covered in Desmin+ muscle fibers) in AT-RvD1 hydrogel treatment as compared to both control hydrogel and untreated controls. (F-M) Second harmonic generation data was recorded within one mean-free-path from the surface of untreated, control hydrogel, AT-RvD1 hydrogel and healthy mouse spinotrapezius muscle. The angular Fourier filter (AFF) histograms are shown in (N-Q). (R) Percentage of muscle fiber area over whole image. Statistical analyses were performed using a one-way analysis of variance (ANOVA) with Tukey’s multiple comparisons; * *p* < 0.05 and ** *p* < 0.01; n = 3 to 4 animals per group. Scale bar = 100 µm.

### 3.8 Dimensionality reduction reveals changes in lipid mediator metabolism after local AT-RvD1 treatment in a quadriceps model of VML injury

The spinotrapezius model is a relatively thin, nearly two dimensional structure, and serves as an initial testbed for therapeutic development for VML. VML injuries in larger muscle groups on the extremities, like the quadriceps, are more challenging to treat than in the spinotrapezius, due to more complex morphological^28^, biomechanical^53^, and micro-circulatory^54,55^ properties. Additionally, injuries in the quadriceps allow for assessment of pain and functional outcomes which cannot be reliably measured in a spinotrapezius injury.

Temporal metabolipidomic profiling of the mouse quadriceps VML was performed with and without local AT-RvD1 treatment. Mice received a 3 mm quadriceps VML injury and were treated with a PEG-4MAL hydrogel that was either unloaded (control hydrogel), or loaded with AT-RvD1 (Fig. 6A). LC-MS/MS lipidomics was performed at 3 and 7 days post-VML. Data were analyzed using partial least squares discriminant analysis (PLSDA) to identify multi-variate patterns and trends in the data that might not be apparent through a more conventional statistical analysis^56,57^.

**Fig. 6:**
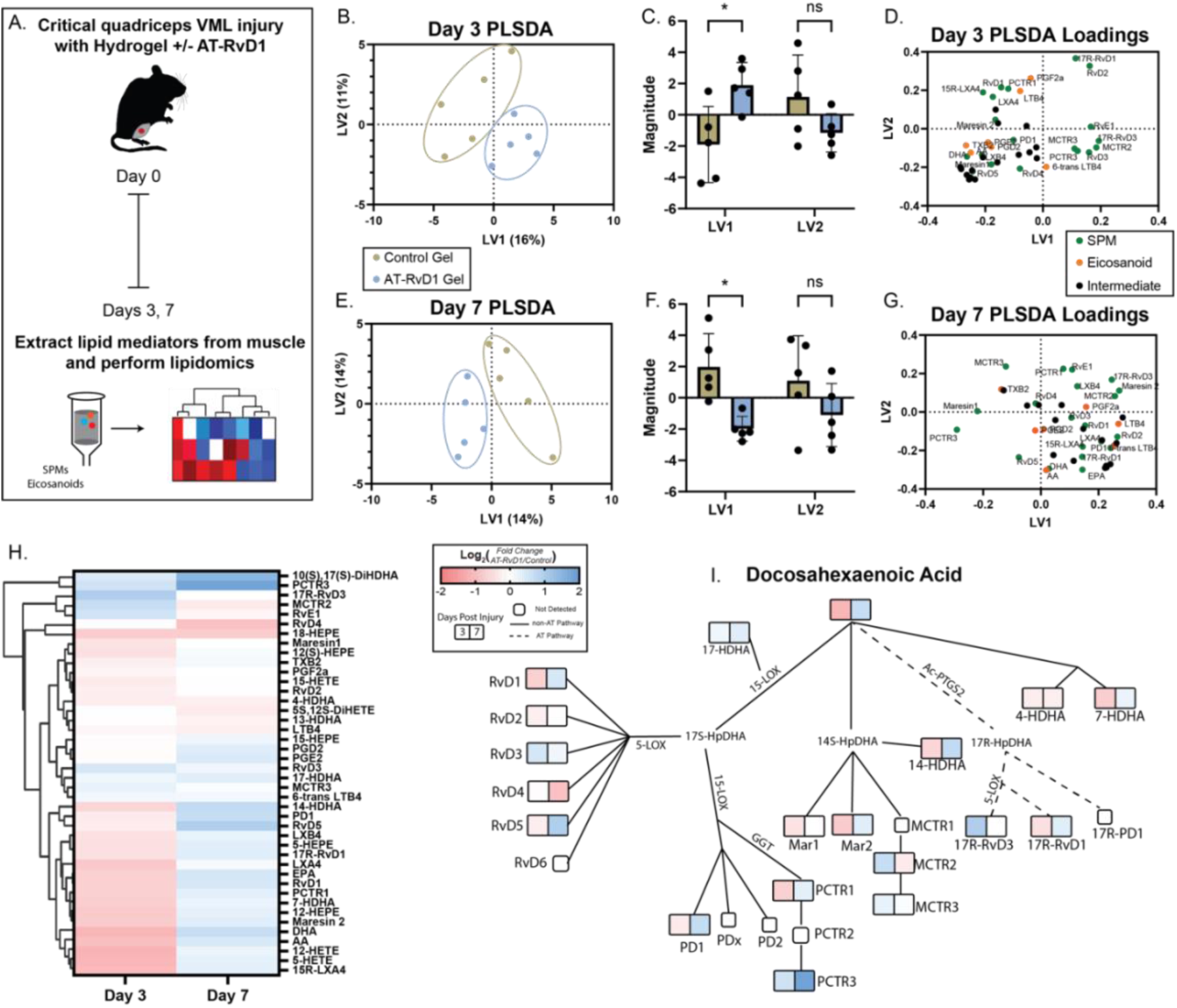
Dimensionality reduction reveals changes in lipid mediator metabolism after local AT-RvD1 treatment in a quadriceps model of VML injury. (A) Experimental outline. All mice received a critical quadriceps VML injury and were treated with either a control hydrogel or a AT-RvD1-loaded hydrogel. Time points were days 3 and 7 for LC-MS/MS lipidomics of the injured quadriceps. PLSDA of lipid mediator concentration in the quadriceps at day 3 (B-D) and day 7 (E-G). (B) PLSDA embedding of control vs AT-RvD1 gel groups at day 3 post-injury. (C) Control and AT-RvD1 gel groups are separated along the LV1 axis at day 3. (D) Loadings plot of each lipid mediators’ contribution to the first two latent variables of the day 3 PLSDA model. (E) PLSDA embedding of control vs AT-RvD1 gel groups at day 7 post-injury. (F) Control and AT-RvD1 gel groups are separated along the LV1 axis at day 7. (G) Loadings plot of each lipid mediators’ contribution to the first two latent variables of the day 7 PLSDA model. (H) Hierarchically clustered heatmap showing average log2 fold change of lipid mediator concentration in the AT-RvD1 vs control gel groups at days 3 and 7 post-injury. (I) Overlay of the heatmap in (H) onto a wireframe diagram of DHA’s metabolic network. For each lipid mediator, the concentration measured at day 3 is represented in the left square, while that at day 7 is represented in the right square. Statistics were a one-way ANOVA between groups. * *p* <0.05. n=5 animals per group.

PLSDA is a supervised dimensionality reduction technique that uses categorical information of samples to find latent variables that best separate the classes or groups. PLSDA is useful for identifying differential features between groups, building classification models, and supervised dimensionality reduction^56,57^. The first latent variable (LV1) explains the most substantial proportion of the variance that discriminates between the control hydrogel and AT-RvD1 hydrogel groups (Fig. 6B), capturing the significant variance between them (Fig. 6C). The loadings plot for LV1 reveals the specific metabolites contributing the most to this separation (Fig. 6D). Variables with high positive or negative loadings on LV1 are the most influential in driving the distinction between the Control and AT-RvD1 hydrogel groups.

The analysis of Day 3 clustering results and PLSDA loadings plot (Fig. 6B-D) provides insights into the lipid mediator profiles of the AT-RvD1 hydrogel group versus the control hydrogel group. Metabolites such as 15R-LXA4, RvD1, LXA4, Maresin 2, DHA, TXB2, LXB4, RvD5, PDG2, PGE2, and PD1 showed negative loadings on LV1, indicating a decrease in these lipid mediators in the AT-RvD1 Gel group compared to the control group. Conversely, positive loadings on LV1 for metabolites such as 17R-RvD1, RvD2, RvE1, 17R-RvD3, MCTR2, RvD3, MCTR3, and PCTR3 indicate an increase in these lipid mediators in the AT-RvD1 Gel group compared to the Control group on Day 3. Negative loadings for Lipoxins (15R-LXA4, LXA4, LXB4) and positive loadings for Resolvins suggest a potential shift from the Lipoxin pathway towards the Resolvin pathway in the AT-RvD1 Gel group.

The analysis of Day 7 clustering results and PLSDA loadings plot (Fig. 6E-G) provide further insight into the lipid mediator profiles of the AT-RvD1 Gel group compared to the Control group. The AT-RvD1 Gel group clustered on the negative LV1 axis and were significantly separated from the Control Gel group, which clustered on the positive LV1 axis (Fig. 6E and F). Metabolites such as PCTR3, Maresin 1, MCTR3, TXB2, and RvD5 showed negative loadings on LV1, indicating a relative increase in the AT-RvD1 Gel group compared to the Control Gel (Fig. 6G). Conversely, metabolites like PCTR1, RvE1, LXB4, 17R-RvD3, Maresin 2, LXA4, PGF2a, RvD3, RvD1, LTB4, RvD2, PD1, 6-trans-LTB4, 17R-RvD1, EPA, 15R-LXA4, PGD2, PGE2, AA, and DHA showed positive loadings on LV1, indicating a relative increase in the Control Gel group at Day 7. Together, the metabolites skewed towards the Control Gel cluster could have heterogenous effects on inflammation resolution, which is consistent with the ambiguous immune phenotypes we have previously observed after untreated murine quadriceps VML^42^. Overall, this shift towards a pro-resolving lipid mediator profile over both timepoints could indicate that local AT-RvD1 hydrogel treatment initiates earlier lipid mediator class-switching, which aligns with the proposed anti-inflammatory and pro-resolution effects of the AT-RvD1-loaded gel.

We observed similar trends in lipid mediator profile when comparing local ("AT-RvD1 Gel”) versus systemic (AT-RvD1 intraperitoneal injection, “IP”) treatments (Fig. S5). At day 3 post-VML injury, the AT-RvD1 Gel group was significantly separated from the IP group on the first latent variable (Fig. S5A and B). The loadings plot for the day 3 comparison suggests that SPMs such as 15R-LXA4, 17R-RvD3, Maresin 2, and PCTR1 were the primary drivers of the AT-RvD1 Gel group clustering on the negative LV1 axis, while pro-inflammatory mediators such as PGF2a, TXB2, PGE2, and PGD2 were the primary drivers of the IP group clustering on the positive LV1 axis (Fig. S5C). These observations, similar to the AT-RvD1 Gel versus Control Gel comparison, indicate earlier lipid mediator class switching occurred with local AT-RvD1 treatment. The AT-RvD1 Gel and IP groups maintained their separation along the LV1 axis 7 days after injury (Fig. S5D and E). The increased importance of the SPMs Maresin 1, 17R-RvD1, and RvD1 on the positive LV1 axis could indicate delayed lipid mediator class-switching with systemic versus local AT-RvD1 administration (Fig. S5F).

The heatmap of Log2 fold change (AT-RvD1/control) in Fig. 6H visually represents the shifts in metabolite levels between Day 3 and Day 7, enabling the qualitative observation of data patterns and trends. Metabolites like RvD3, MCTR3, and PCTR3 showed increased levels (blue) at both Day 3 and Day 7 in the AT-RvD1 Gel group compared to the Control group. This is consistent with earlier loading plot observations, confirming the sustained biosynthesis of these pro-resolving and anti-inflammatory SPMs induced by the AT-RvD1-containing gel. Certain pro-inflammatory eicosanoids like LTB4 and TXB2 showed decreased levels (red) at both time points in the AT-RvD1 Gel group, which aligns with the proposed anti-inflammatory effects of the Resolvin formulation. DHA and EPA, which are precursors for SPM biosynthesis, have increased levels (blue) at Day 7, indicating their enhanced metabolism to produce SPMs at this later time point.

Several metabolites, including PD1, RvD5, and Maresin 2, shifted from decreased levels (red) at Day 3 to increased levels (blue) at Day 7 in the AT-RvD1 Gel group, suggesting a temporal regulation of different SPM pathways or a potential feedback mechanism. The heatmap depicted in Fig. S6A enables a comparison of metabolite levels between local and systemic AT-RvD1 administration at Day 3 and Day 7. The heightened levels of multiple SPMs at Day 3 in the AT-RvD1 Gel group, coupled with the upregulation of DHA and EPA, indicate an earlier onset of SPM production facilitated by local AT-RvD1 delivery, contrasting with the delayed elevation of SPM levels observed at Day 7 in the IP group. Overall, the heatmap provides a comprehensive view of the metabolic shifts induced by the AT-RvD1-containing gel, supporting the proposed pro-resolving and anti-inflammatory effects observed in previous analyses. It also emphasizes the complex temporal regulation of different lipid mediator pathways and their potential interactions during the resolution of inflammation.

Fig. 6I, Fig. S6B-D, and Fig. S7 present network diagrams illustrating the metabolic pathways of various metabolites involved in the study. These diagrams were simplified from the Reactome.org diagram viewer^58^, grouping metabolites into the distinct clusters of D-series Resolvins, Maresins, and Protectins (Fig. 6I and S6D); Prostaglandins, Leukotrienes, and Lipoxins (Fig. S6B and S7A); and E-series Resolvins (Fig. S6C and S7B). By comparing these network diagrams with the heatmaps in Fig. 6H and Fig. S6A, we can infer the temporal dynamics of metabolic pathways over time. These findings support the hypothesis that local delivery of the AT-RvD1-containing biomaterial modulates the inflammatory response. This promotes a pro-resolving lipid mediator profile which aids muscle regeneration after a VML injury. The data provide insights into the specific metabolic pathways and temporal dynamics involved in this process, underscoring the potential therapeutic benefits of the AT-RvD1-containing gel formulation.

### 3.9 Local AT-RvD1 delivery in quadriceps VML results in improved muscle regeneration

By 14 days post-VML in the quadriceps model, there remained a large area devoid of myofibers with pockets of fatty infiltration. In addition to the large defect region in critical VML injuries, there was also an increase in highly cellularized areas in between individual myofibers, characteristic of unresolved inflammation^28^. In these studies, we compared vehicle control hydrogel and AT-RvD1-containing hydrogel treated non-healing VML injuries in the murine quadriceps (Fig. 7A). We evaluated whether the local hydrogel delivery of AT-RvD1 ameliorates many of the hallmark human clinical features that are recapitulated by our pre-clinical model and results in facilitated muscle regeneration (Fig. 7B).

**Fig. 7:**
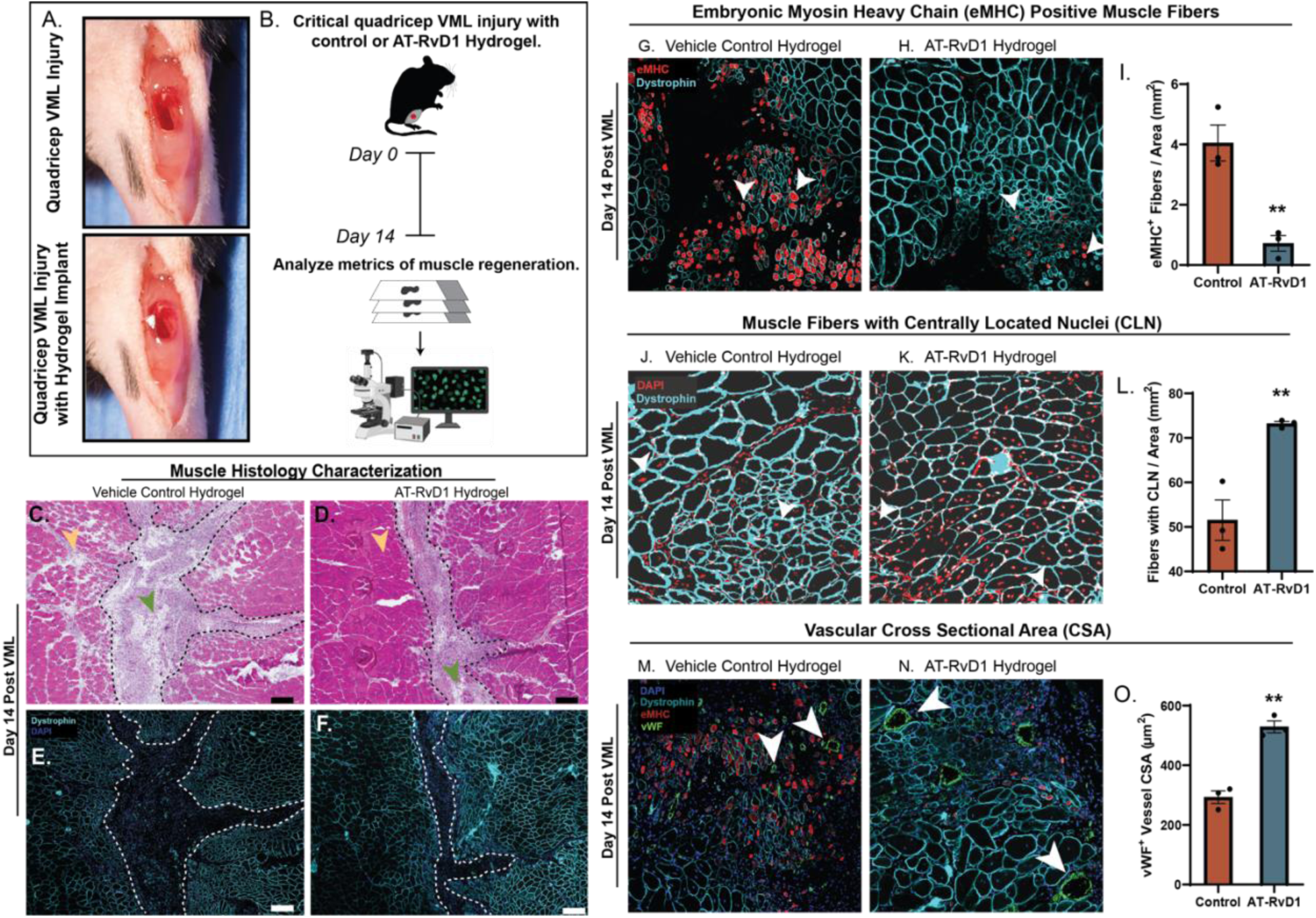
Pro-resolving hydrogel treatment to a quadriceps VML model induces a regulated muscle regeneration process. (A) Critical VML injury was performed in a murine quadriceps model. An unloaded (control) or AT-RvD1-loaded hydrogel was prepared for in situ gelation into the defect area. (B) Experimental schematic of outcome measures at 14 days post-VML. Representative H&E stains (C and D) and corresponding IHC stains (E and F) of quadriceps cross sections 14 days post-VML. Myofibers identified with dystrophin (cyan) and cell nuclei identified via DAPI (blue). Representative images of the defect area showing eMHC^+^ myofibers (white arrows) with eMHC expression (red) and a ring of dystrophin (cyan) in the vehicle control and pro-resolving hydrogel groups (G and H). Quantification of eMHC^+^ fibers normalized to area (I). Representative images of the defect area showing centrally located nuclei as myofibers (white arrows), seen as a dystrophin^+^ (cyan) ring with a centrally located nuclei (red) (J and K). Quantification of number of centrally located nuclei between vehicle control and AT-RvD1 groups normalized to area (L). Representative images of vessels (white arrows), represented by a ring of endothelial cells (vWF^+^; green), around regenerating myofibers in vehicle control (M) and pro-resolving hydrogel (N) treated VML injuries 14 days post-VML. Quantification of the average cross-sectional area of vessels (O). Statistical analyses were performed using two-tailed *t* tests; ** *p* < 0.01; n = 3 animals per group (I, L, O).

Cross-sections of mouse quadriceps at 14 days post-VML were stained with hematoxylin and eosin (H&E) and immunohistochemically stained for dystrophin and DAPI to assess gross muscle healing after local AT-RvD1 treatment compared to vehicle control (Fig. 7C-F). We observed a considerable decrease in the size of the injury area with pro-resolving hydrogel treatment compared to control (Fig. 7C-F). Many of the hallmark VML pathological features, which were evident in the vehicle control group, were not present in injuries treated with local AT-RvD1, as little to no areas were devoid of myofibers, no irregular myofiber organization, no increase in area of non-muscle cell types between myofibers (yellow arrow) and there was no indication of fatty infiltration (green arrow) (Fig. 7C and D).

At 14 days post-VML, we assessed muscle regeneration in the defects by quantifying myofibers which were embryonic myosin heavy chain (eMHC) positive or had centrally located nuclei (Fig. 7G-L). eMHC is considered a marker for newly regenerated myofibers as it is transiently expressed during the early stages of myofiber development, considered to be in the first 7 days post-VML^59^. However, the persistent expression of eMHC at later stages of regeneration indicates a continuous cycle of failed attempts at myofiber regeneration. On the other hand, centrally located nuclei, another marker for regenerating myofibers, persists longer than eMHC to indicate the presence of regenerating fibers that were able to mature. Representative images of the defect area between vehicle control and AT-RvD1 hydrogel show a striking difference in number of eMHC^+^ myofibers (Fig. 7G and H; white arrows). This was quantified as number of eMHC^+^ myofibers normalized to area, and our results showed a 4-fold decrease in the presence of this early marker of regenerating myofibers with AT-RvD1 treatment (Fig. 7I). Conversely, there was a 50% increase in the number of myofibers with centrally located nuclei (Fig. 7J and K; white arrows) in the defect area with local AT-RvD1 delivery at 14 days post-VML injury (Fig. 7L). This increase in a marker of myofiber regeneration that was sustained in successful regeneration indicates that local AT-RvD1 delivery promotes muscle regeneration post-VML.

A key component in successful muscle regeneration is also the successful formation of a vascular network after injury. Thus, we visualized endothelial cells by staining for von Willebrand Factor (vWF) and identified vessels within the defect area and around regenerating fibers (Fig. 7M and N). We quantified the average cross-sectional area of vessels within the defect area and observed a roughly 40% increase in the size of newly formed vessels with local AT-RvD1 treatment at 14 days post-VML injury (Fig. 7O). This indicates that regenerating muscle with AT-RvD1 treatment 14 days post-VML is more vascularized compared to control hydrogel treatment.

These results show that local AT-RvD1 delivery in a critical-sized quadriceps VML model decreased the area of the defect and resulted in a more compact myofiber structure around the defect at 14 days post-VML (Fig. 7C-F). This improvement in muscle healing was quantified via the significant decrease in an early marker of muscle regeneration (eMHC) and the increase of markers of more mature regenerating myofibers (centrally located nuclei) (Fig. 7G-L). Furthermore, there was a significant increase in the area of vessels surrounding the regenerating myofibers with AT-RvD1 treatment (Fig. 7M-O). Taken together, these results show that hydrogel-mediated delivery of AT-RvD1 promoted the successful regeneration of muscle after a non-healing quadriceps VML injury.

### 3.10 Local AT-RvD1 delivery improves function and reduces pain after quadriceps VML

The main clinical challenge in treating traumatic VML injuries is the recovery of muscle function after treatment. This inability to recover function following such an injury often results in permanent disability and chronic and pathologic pain^2,60–62^. Thus, for clinical translation, two important endpoints of a VML therapeutic are the quantification of *in vivo* force generation and reduction of pain after injury. This pre-clinical quadriceps VML injury presented with a sustained loss of function similar to that of clinical VML injuries. Having established the ability of our pro-resolving hydrogel therapeutic to increase biosynthesis of SPMs, promote the cellular resolution of inflammation, and increase various metrics of muscle regeneration, we tested the ability of this therapeutic to recover muscle function and reduce associated chronic pain (Fig. 8A).

**Fig. 8:**
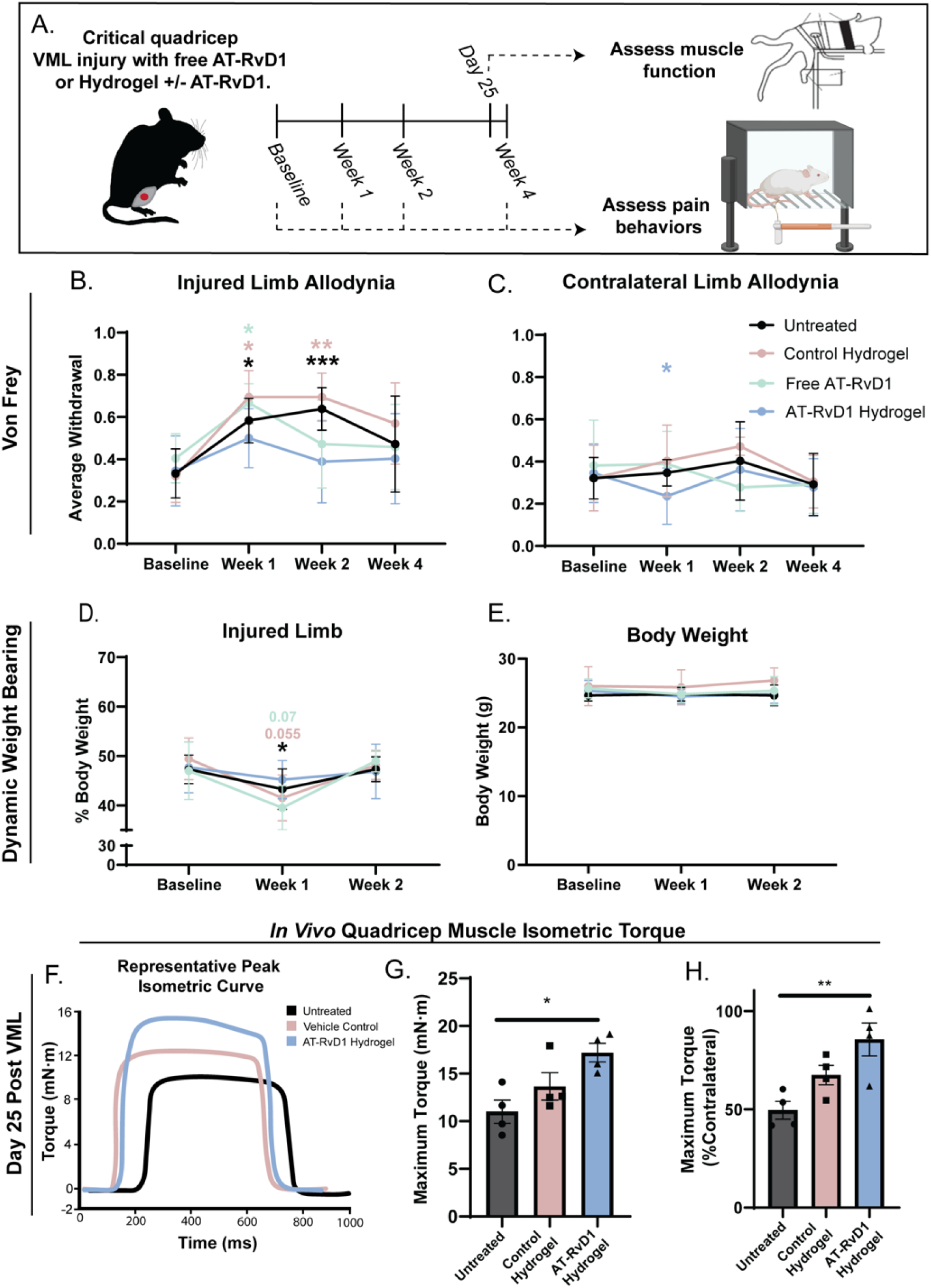
Local AT-RvD1 delivery after quadriceps VML results in a reduction in pathologic pain and an improvement in muscle function. (A) Experimental schematic. Baseline pain behaviors were assessed by Von Frey before VML surgery. Mice then received a critically-sized quadriceps VML injury and received either no treatment, a local injection of free AT-RvD1 in PBS, or a hydrogel implant with or without AT-RvD1. Pain behaviors were assessed at weeks 1, 2, and 4 post-VML. In a separate experiment, mice received a critically-sized VML injury, received either no treatment, a control hydrogel, or a AT-RvD1-loaded hydrogel, then their quadriceps function was assessed via isometric tetanic torque at day 25. (B) and (C) Mechanical allodynia of the injured (B) and contralateral (C) limbs was assessed using a series of VF filaments. A higher average withdrawal indicates elevated pain. (D) Spontaneous pain behaviors were assessed using DWB by measuring the percentage of body weight placed on the injured limb. Week 4 measurements were not acquired due to sensor malfunction. (E) Body mass measurements at each timepoint throughout the study period. (F-H) Quadriceps function was assessed by measuring the maximal isometric tetanic torque about the knee after electrical stimulation of the femoral nerve. Smoothed representative isometric force curves for each group (F) shows a sustained force production during quadricep stimulation. The maximum torque was quantified and reported as mN-m (G). This torque was normalized to the torque produced from the healthy contralateral muscle to quantify the percentage of force generated in the injured quad compared to a healthy muscle (H). Statistical analyses for B-E were performed using a two-way ANOVA with Holm-Sidak’s *post-hoc* test for comparing post-op timepoints vs. baseline within each group. Stars are color-coded by group to represent significant differences from that group’s baseline. n=6 animals per group (B-E). Statistical analyses for G and H were one-way ANOVA with Tukey’s multiple comparisons. n=4 animals per group (G-H). * *p* <0.05, ** *p* <0.01, *** *p* <0.001.

Fig. 8B-H presents the impact of AT-RvD1 hydrogel treatment in a preclinical quadriceps VML model, highlighting its benefits in enhancing muscle function and alleviating pathological pain. At 1-and 2-weeks post-injury, animals treated with AT-RvD1 hydrogel did not exhibit elevated hypersensitivity-associated behaviors, whereas untreated and control groups did. This is evidenced by withdrawal responses to benign mechanical stimuli in the AT-RvD1 hydrogel-treated group which did not significantly deviate from baseline (Fig. 8B). Resolvins are known to exert potent analgesic effects, likely due to their anti-inflammatory and pro-resolving actions. Furthermore, this early reduction in mechanical allodynia underscores the hydrogel’s ability to influence pain processing mechanisms that develop during the acute post-injury phase^63–66^, bolstering its therapeutic potential for managing injury-induced hypersensitivity. Continued alleviation of mechanical allodynia into Week 2 supports the sustained efficacy of the AT-RvD1 treatment. Notably, a significant decrease in mechanical hypersensitivity was observed in the contralateral limb of the AT-RvD1 gel-treated animals at Week 1, suggesting a potential modulation of central sensitization processes (Fig. 8C).

Dynamic weight-bearing assessments (Fig. 8D) at Weeks 1 and 2 further validated these findings, with AT-RvD1 gel-treated animals showing no change in weight bearing on the injured limb, indicative of diminished spontaneous pain and enhanced functional limb use. These results correlated with the reductions in mechanical allodynia, affirming the dual benefits of the treatment. Consistent body masses across all groups throughout the study period (Fig. 8E) indicate that the treatments did not adversely affect overall animal health and growth. Together, the dynamic weight-bearing and von Frey assessments substantiate the therapeutic efficacy of AT-RvD1 hydrogel in mitigating pain and facilitating functional recovery after muscle injury^40^.

We next assessed the ability of the pro-resolving hydrogel therapeutic to improve muscle function at 25 days post-VML injury compared to untreated and vehicle control hydrogel treated VML injuries. We first compared the maximal torque produced between untreated, vehicle control hydrogel, and AT-RvD1 hydrogel-treated VML injuries (Fig. 8F). Muscles treated with AT-RvD1 hydrogels produced roughly 75% more torque than an untreated VML injury at 25 days post-VML (Fig. 8G). Furthermore, we normalized the force produced in each injured quadriceps to its corresponding healthy contralateral quadriceps muscle. This produced a percentage indicative of how similar the injured quadriceps performs to the healthy control in terms of function (e.g., 100% indicated complete functional equivalence with the uninjured contralateral muscle). Using this metric, the untreated VML muscle was only able to produce, on average, 50% of the force compared to its contralateral control (Fig. 8H). Strikingly at 25 days post-injury, VML quadriceps muscle treated with the pro-resolving hydrogel restored 86% of the force compared to its contralateral control, a significant increase compared to untreated VML injuries (Fig. 8H). These data confirm that local hydrogel-mediated AT-RvD1 delivery is an effective way to restore muscle function after critical VML injury.

## 4. DISCUSSION

Volumetric muscle loss (VML) injuries are a significant healthcare burden that can arise from various traumatic and non-traumatic scenarios, including traumatic injuries, congenital birth defects, and tumor resection. Current clinical strategies for VML treatment, such as muscle flap autografts and free tissue transfer, have limited success in promoting functional improvement and patient quality of life^1,2^. While regeneration following VML injury is similar to epimorphic regeneration, in which some vertebrates regrow complete functional appendages after previous amputation, the formation of a blastema-like structure is not observed in muscle regeneration. Moreover, muscle stem cells, in addition to being largely ablated in traumatic muscle injuries, must overcome the negative impact of the microenvironment. Cytokine imbalances, immune cell infiltration, and fibrosis create a hostile environment for muscle regeneration^2,3^, and persistent inflammatory cells, including neutrophils and NK cells, produce reactive oxygen species (ROS) that disrupt the microenvironment required for muscle stem cell proliferation and differentiation^3^. Limited success of bioactive materials, such as decellularized skeletal muscle and porcine small intestine submucosa (SIS-ECM), has been reported in preclinical models^67–69^, and cell-based therapies, including delivery of MuSCs, have been shown to promote newly formed aligned functional fibers in thin and partial thickness preclinical VML defect models^70–72^. However, the persistent inflammation of large-volume skeletal muscle can overwhelm myogenic cells, often leading to fibrosis and strength deficits. Therefore, there is a need for targeted therapies that can promote the resolution of inflammation while also stimulating repair and regeneration after VML.

Our study utilized mass spectrometry-based lipidomics to measure lipid mediators derived from AA, DHA, and EPA in a pre-clinical VML model. Similar studies in models of acute muscle injury and functional regeneration, show a successful lipid mediator class switch that results in the production of downstream SPMs^19^. Additionally, a recent study in the tibialis anterior VML model suggests that such an injury is characterized by a sustained pro-inflammatory lipid mediator response^73^. However, our results suggest a robust but transitory pro-inflammatory response, characterized by a sharp spike in Prostaglandins at 1 day post-VML that return to uninjured baseline by 3 days post-VML. The lack of downstream, bioactive pro-resolving lipid metabolites detected in the methods used to characterize the tibialis anterior VML injury offered a partial view of the lipid mediator response to VML injury^73^. However, given our ability to robustly detect downstream SPMs of various fatty acid origins, we were able to reveal a unique impairment in the ability of the local injury environment to produce SPMs after a VML injury. An observation evidenced by the lack of accumulation of SPMs derived from AA, DHA and EPA.

Importantly, the availability of AA, DHA, and EPA to act as a substrate for downstream lipid mediator synthesis was not the rate-limiting step in this process as we see a significant accumulation of these fatty acids after VML injury. This implies that it is the enzymatic reactions downstream of the fatty acids that may be dysregulated following VML injury. Multiple enzyme isoforms that participate in lipid mediator production exist, which differ in their substrate specificity, cell expression profiles, and enzyme kinetics. Additionally, the production of SPMs often involves transcellular biosynthetic routes involving multiple enzymes. How the activity of these enzymes is affected in traumatic and chronic injuries along with a spatial analysis of how cellular interactions are affected after the large ablation of skeletal muscle is a focus of future studies.

Our findings suggest that the local injury environment in VML may impede the production of SPMs and contribute to the limited success of current VML treatment strategies. Therefore, we designed a strategy to promote the lipid mediator switch that produces SPMs during inflammation.

This strategy focused on promoting the initial activation of the local pro-resolving signaling pathways that would serve to limit the duration of the downstream inflammatory cascade. Successful resolution of acute inflammation is self-limiting and requires the active regulation of the inflammatory process to re-establish homeostasis. Our approach is based on the hypothesis that releasing AT-RvD1 within the first 24 hours of the inflammatory injury via synthetic hydrogels will rebalance inflammatory mediator metabolism and create a pro-regenerative microenvironment for muscle repair.

Our studies have shown that synthetic PEG-4MAL hydrogels that can encapsulate and locally deliver the SPM Aspirin-triggered Resolvin D1 (AT-RvD1) to shift the local immune cell milieu towards a pro-regenerative phenotype^26^. Hydrogels can be injected directly into the tissue, where it undergoes gelation under physiological conditions and firmly integrates with biological tissues due to the maleimide functional groups reacting with thiol groups present in the tissue^74^. Our data show that we successfully promote the production and accumulation of SPMs at 72 hours post-VML with local AT-RvD1 treatment beyond the window of exogenous AT-RvD1 release. In our analysis of spinotrapezius VML, we observed a broad spectrum of SPM classes being synthesized, which include multiple metabolites in the DHA pathway as well as other SPMs in the AA and EPA pathways. Further analysis in the quadriceps VML model underscores the efficacy of AT-RvD1 in modulating the lipid mediator landscape post-VML injury, highlighting not only a reduction in pro-inflammatory mediators but also an enhancement of pro-resolving ones. This dual effect is central to the improved outcomes observed, indicating a significant shift towards a pro-regenerative environment following treatment.

Our results also highlight AT-RvD1’s ability to limit neutrophil infiltration after local delivery. Specifically, our analysis shows that local delivery of AT-RvD1 reduced the total concentration of neutrophils in the tissue and limited the polarization of neutrophils to the aged phenotype. Because the aged neutrophil phenotype is known to have a greater propensity to undergo NETosis and display elevated expression of MPO, reducing infiltration of overall neutrophils and pro-inflammatory subsets could be important to the anti-fibrotic responses to AT-RvD1 hydrogels^75^. The function of local AT-RvD1 to limit pro-inflammatory and pro-fibrotic cell polarization is also reflected in the shift in the macrophage population towards a pro-resolving and pro-regenerative M2 phenotype. Lipid mediator profiling also correlates with the observed shifts in neutrophil and macrophage phenotypes, suggesting the increased levels of SPMs foster a milieu that favors the M2 macrophage phenotype and limits neutrophil infiltration and polarization, thereby aiding in muscle regeneration and reducing fibrosis. As such, our approach promotes the resolution of inflammation at both the molecular and cellular levels.

These findings also suggest that activating pro-resolving signaling pathways with local delivery of exogenous pro-resolution cues during the onset of inflammation enhances the regenerative capacity of the skeletal muscle after VML injury. Classical ‘anti-inflammatory’ strategies to reduce chronic inflammation associated with VML have been proposed, but significant concerns persist about the undesired impairment of tissue regeneration^14,76^. As the spinotrapezius muscle has the ability to be whole mounted and imaged throughout the full thickness of the muscle, we leveraged this feature of the model and used whole-mount IHC and SHG imaging to demonstrate the ability of our local therapeutic to improve muscle healing after VML significantly. The combination of these two imaging methods shows that AT-RvD1 hydrogels implanted into a VML injury results in near complete defect closure after 14 days and a compact muscle fiber structure with a mature and organized collagen network. Delivery of AT-RvD1 from PEG-4MAL hydrogels also increases the pool of muscle progenitor cells – FAPs and MuSCs – that are essential to forming new ECM and muscle fibers, respectively, after a muscle injury. The direct signaling interaction of Resolvins with MuSCs has been previously explored^19^; however, their direct signaling interaction on FAPs is poorly understood. We show that AT-RvD1 can directly influence the fate of FAP differentiation by decreasing their propensity for fibrotic differentiation. This further illustrates the important dual nature of AT-RvD1 to act on both immune cells to promote resolution and on muscle progenitor cells to promote muscle regeneration and limit fibrosis.

In the pre-clinical quadriceps VML model, our pro-resolving hydrogel treatment via local AT-RvD1 delivery shifted the lipid mediator profile towards a pro-resolving one and significantly improved muscle healing outcomes. Our results demonstrated that at 25 days post-injury, VML injuries treated with the pro-resolving hydrogel generated 86% of the force of a healthy muscle, whereas untreated injuries only generated 50% of the force. This functional recovery was achieved via the biomaterial mediated delivery of one dose of an SPM. Furthermore, a therapeutic that ameliorates the additional clinical challenge of chronic pain is vital to the complete recovery of a patient with a traumatic injury. Interestingly, our VF and DWB results suggest that this single dose of SPM has sustained analgesic effects. In addition, the treatment reduced the total defect area at 14 days post-VML injury and reduced the presence of non-muscle cell types between regenerated myofibers. While control hydrogel-treated, VML injuries exhibited delayed healing, possibly due to continuous attempts and disruptions of muscle regeneration, treatment AT-RvD1 hydrogels ameliorated this response by reducing the expression of eMHC and increasing the number of mature regenerated fibers. Additionally, the treatment significantly increased the vascular area around regenerating fibers in the defect area at 14 days post-VML. These promising results demonstrate the significant potential of our pro-resolving hydrogel treatment to improve the treatment of VML injuries, highlighting its translational potential for future functional studies and traumatic injuries.

The importance of a local and biomaterial-driven therapeutic approach is underscored by comparing it with systemic or biomaterial-free delivery of exogenous SPMs. Recent studies, including those by Giannakis et al., Markworth et al., and Dort et al.^17–19^, have demonstrated functional improvements or increased regeneration with injection of exogenous RvD2 or RvD1. More recently, the study by Castor-Macias et al. investigated the effects of Maresin 1 (MaR1) treatment on muscle regeneration and functional recovery after VML injury in mice, showing statistically significant improvements in maximal tetanic force production of the tibialis anterior (TA) muscle^73^. However, this was achieved through intramuscular injection every two days without a biomaterial delivery vehicle, and muscle strength testing was performed via direct electrical muscle stimulation. Our findings, however, indicate that a single local dose delivered by a hydrogel significantly enhances both muscle regeneration and functional recovery. Using femoral nerve stimulation for muscle strength testing, we observed that the injured quadriceps’ strength was restored to as much as 86% of that of uninjured contralateral quadriceps. The distinction between stimulation methods is crucial, as nerve stimulation provides a more physiologically-relevant assessment of muscle function post-VML injury by accounting for muscle innervation. Thus, our study demonstrated a more clinically-relevant restoration of muscle strength following the administration of SPMs in a biomaterial.

The biotech/pharmaceutical field is reaching a stage in which the importance of therapeutics that activate resolution is coming into focus. This is marked by the progression of several synthetic, small molecule, and peptide agonists of the RvD1 receptor FPR2 into clinical trials (NCT05372107, NCT03335553, TrialTroveID-491584), a significant achievement in the field of resolution pharmacology. In addition to small molecule and peptide agonists of pro-resolving receptors, lipid mediator mimetics are also being progressed into clinical trials^77^. Furthermore, a few groups have investigated local delivery of SPMs^78–81^, which support our findings that localized delivery strategies are beneficial for enhancing the functional efficacy of pro-resolving therapeutics. While our studies are constrained to the use of the endogenous lipid AT-RvD1 as a therapeutic, future studies will evaluate the potential increase in efficacy of alternative pro-resolving molecules when locally delivered via our engineered platform. Additionally, while we focus on a traumatic injury in this study; in future studies, our biomaterial-based pro-resolution therapeutic strategy will be evaluated in other models of chronic inflammation such as those being pursued in the clinic with synthetic pro-resolving molecules (i.e. Ulcerative Colitis, Chron’s Disease, Psioriasis, etc.).

In summary, this study demonstrates that VML injuries are characterized by a deficiency in the biosynthesis of pro-resolving lipid mediators and a persistent dysregulated inflammatory response. We show that biomaterial-mediated delivery of AT-RvD1 can locally resolve persistent inflammation and promote a pro-regenerative cellular environment, resulting in significantly improved muscle healing outcomes. Additionally, we have previously shown that delivering MuSCs in this PEG-4MAL hydrogel system supports their survival, proliferation, and differentiation, while also enhancing their engraftment into cryo-injured muscles of dystrophic and aged mice compared to gel-free and collagen gel controls^74^. These findings suggest that our innovative pro-resolving platform holds great promise for combining with stem cell therapy to treat traumatic skeletal muscle injuries. This work sets the stage for future development and scale-up of hydrogel-based therapy for functional studies and clinical translation.

## AUTHOR INFORMATION

### Author Contributions

T.C.T., F.S.P., H.Z., M.S., P.Q., A.J.G, L.J.M, Y.C.J, N.J.W, and E.A.B designed the research, analyzed the data and wrote the manuscript. T.C.T., F.S.P., H.Z., L.A.H., T.Z., M.B., S.E.A., J.A.H., K.A.L., M.A.A., K.M-S., X.L., X.Y., and H.S.L. performed the research, analyzed the data, and reviewed the manuscript. All authors have given approval to the final version of the manuscript.

## ACKNOWLEDGEMENT

We thank the core facilities at the Parker H. Petit Institute for Bioengineering and Bioscience at the Georgia Institute of Technology and at the Emory Integrated Lipidomics core for the use of their shared equipment, services, and expertise. Graphical abstract muscle images and experimental timeline schematics were created with BioRender.com.

## REFERENCES

1 Garg, K. et al. Volumetric muscle loss: persistent functional deficits beyond frank loss of tissue. Journal of Orthopaedic Research 33, 40–46 (2015).

2 Corona, B. T., Rivera, J. C., Owens, J. G., Wenke, J. C. & Rathbone, C. R. Volumetric muscle loss leads to permanent disability following extremity trauma. Journal of Rehabilitation Research & Development 52 (2015).

3 Larouche, J., Greising, S. M., Corona, B. T. & Aguilar, C. A. Robust inflammatory and fibrotic signaling following volumetric muscle loss: a barrier to muscle regeneration. Cell death & disease 9, 409 (2018).

4 Goldring, M. B. & Otero, M. Inflammation in osteoarthritis. Current opinion in rheumatology 23, 471–478 (2011).

5 Carnes, M. E. & Pins, G. D. Skeletal muscle tissue engineering: biomaterials-based strategies for the treatment of volumetric muscle loss. Bioengineering 7, 85 (2020).

6 Das, S. et al. Pre-innervated tissue-engineered muscle promotes a pro-regenerative microenvironment following volumetric muscle loss. Communications biology 3, 330 (2020).

7 Chen, Z., Bozec, A., Ramming, A. & Schett, G. Anti-inflammatory and immune-regulatory cytokines in rheumatoid arthritis. Nature Reviews Rheumatology 15, 9–17 (2019).

8 Abramson, S. & Amin, A. Blocking the effects of IL-1 in rheumatoid arthritis protects bone and cartilage. Rheumatology 41, 972–980 (2002).

9 Funk, C. D. Prostaglandins and leukotrienes: advances in eicosanoid biology. science 294, 1871–1875 (2001).

10 Su, W.-H. et al. Nonsteroidal anti-inflammatory drugs for wounds: pain relief or excessive scar formation? Mediators of inflammation 2010 (2010).

11 Tarnawski, A. S. & Jones, M. K. Inhibition of angiogenesis by NSAIDs: molecular mechanisms and clinical implications. Journal of molecular medicine 81, 627–636 (2003).

12 Buckley, C. D., Gilroy, D. W. & Serhan, C. N. Proresolving lipid mediators and mechanisms in the resolution of acute inflammation. Immunity 40, 315–327 (2014).

13 Serhan, C. N. Pro-resolving lipid mediators are leads for resolution physiology. Nature 510, 92–101 (2014).

14 Serhan, C. N. & Savill, J. Resolution of inflammation: the beginning programs the end. Nature immunology 6, 1191–1197 (2005).

15 Bannenberg, G. L. et al. Molecular circuits of resolution: formation and actions of resolvins and protectins. The Journal of Immunology 174, 4345–4355 (2005).

16 Markworth, J. F. et al. Metabolipidomic profiling reveals an age-related deficiency of skeletal muscle pro -resolving mediators that contributes to maladaptive tissue remodeling. Aging Cell 20, e13393 (2021).

17 Giannakis, N. et al. Dynamic changes to lipid mediators support transitions among macrophage subtypes during muscle regeneration. Nature immunology 20, 626–636 (2019).

18 Dort, J. et al. Resolvin-D2 targets myogenic cells and improves muscle regeneration in Duchenne muscular dystrophy. Nature Communications 12, 6264 (2021).

19 Markworth, J. F., et al. Resolvin D1 supports skeletal myofiber regeneration via actions on myeloid and muscle stem cells. JCI insight 5 (2020).

20 Sun, A. R. et al. Pro-resolving lipid mediator ameliorates obesity induced osteoarthritis by regulating synovial macrophage polarisation. Scientific reports 9, 426 (2019).

21 Habouri, L., et al. Deletion of 12/15-lipoxygenase accelerates the development of aging-associated and instability-induced osteoarthritis. Osteoarthritis and cartilage 25, 1719–1728 (2017).

22 Huang, J. et al. Targeting the D series resolvin receptor system for the treatment of osteoarthritis pain. Arthritis & Rheumatology 69, 996–1008 (2017).

23 Juban, G. & Chazaud, B. Efferocytosis during skeletal muscle regeneration. Cells 10, 3267 (2021).

24 Hong, S. et al. Resolvin E1 metabolome in local inactivation during inflammation-resolution. The Journal of Immunology 180, 3512–3519 (2008).

25 Arita, M. et al. Metabolic inactivation of resolvin E1 and stabilization of its anti-inflammatory actions. Journal of biological chemistry 281, 22847–22854 (2006).

26 Sok, M. C. P. et al. Dual delivery of IL-10 and AT-RvD1 from PEG hydrogels polarize immune cells towards pro-regenerative phenotypes. Biomaterials 268, 120475 (2021).

27 Phelps, E. A. et al. Maleimide cross-linked bioactive PEG hydrogel exhibits improved reaction kinetics and cross-linking for cell encapsulation and in-situ delivery. *Advanced materials (Deerfield Beach*, Fla.) 24, 64 (2012).

28 Anderson, S. E. et al. Determination of a critical size threshold for volumetric muscle loss in the mouse quadriceps. Tissue Engineering Part C: Methods 25, 59–70 (2019).

29 San Emeterio, C. L., et al. Nanofiber-based delivery of bioactive lipids promotes pro-regenerative inflammation and enhances muscle fiber growth after volumetric muscle loss. Frontiers in Bioengineering and Biotechnology 9, 650289 (2021).

30 Dalli, J., Colas, R. A., Walker, M. E. & Serhan, C. N. Lipid mediator metabolomics via LC-MS/MS profiling and analysis. Clinical Metabolomics: Methods and Protocols, 59–72 (2018).

31 Qiu, P. et al. Extracting a cellular hierarchy from high-dimensional cytometry data with SPADE. Nature biotechnology 29, 886–891 (2011).

32 Zheng, T. et al. Imaging mitochondria through bone in live mice using two-photon fluorescence microscopy with adaptive optics. Frontiers in Neuroimaging 2, 959601 (2023).

33 Forouhesh Tehrani, K., Pendleton, E. G., Southern, W. M., Call, J. A. & Mortensen, L. J. Spatial frequency metrics for analysis of microscopic images of musculoskeletal tissues. Connective tissue research 62, 4–14 (2021).

34 Quadros, A. U. et al. Dynamic weight bearing is an efficient and predictable method for evaluation of arthritic nociception and its pathophysiological mechanisms in mice. Scientific reports 5, 14648 (2015).

35 Dent, J. O. et al. Advanced dynamic weight bearing as an observer-independent measure of hyperacute hypersensitivity in mice. Canadian Journal of Pain 7, 2249060 (2023).

36 Xu, Z.-Z. et al. Inhibition of mechanical allodynia in neuropathic pain by TLR5-mediated A-fiber blockade. Nature medicine 21, 1326–1331 (2015).

37 Lolignier, S., Eijkelkamp, N. & Wood, J. N. Mechanical allodynia. Pflügers Archiv-European Journal of Physiology 467, 133–139 (2015).

38 Dhandapani, R. et al. Control of mechanical pain hypersensitivity in mice through ligand-targeted photoablation of TrkB-positive sensory neurons. Nature communications 9, 1640 (2018).

39 Chaplan, S. R., Bach, F. W., Pogrel, J., Chung, J. & Yaksh, T. Quantitative assessment of tactile allodynia in the rat paw. Journal of neuroscience methods 53, 55–63 (1994).

40 Li, M. T. A., Willett, N. J., Uhrig, B. A., Guldberg, R. E. & Warren, G. L. Functional analysis of limb recovery following autograft treatment of volumetric muscle loss in the quadriceps femoris. Journal of biomechanics 47, 2013–2021 (2014).

41 Pedregosa, F. et al. Scikit-learn: Machine learning in Python. the Journal of machine Learning research 12, 2825–2830 (2011).

42 Hymel, L. A. et al. Identifying dysregulated immune cell subsets following volumetric muscle loss with pseudo-time trajectories. Communications Biology 6, 749 (2023).

43 Han, W. M., Mohiuddin, M., Anderson, S. E., García, A. J. & Jang, Y. C. Co-delivery of Wnt7a and muscle stem cells using synthetic bioadhesive hydrogel enhances murine muscle regeneration and cell migration during engraftment. Acta biomaterialia 94, 243–252 (2019).

44 San Emeterio, C. L., Olingy, C. E., Chu, Y. & Botchwey, E. A. Selective recruitment of non-classical monocytes promotes skeletal muscle repair. Biomaterials 117, 32–43 (2017).

45 Hymel, L. A. et al. Modulating local S1P receptor signaling as a regenerative immunotherapy after volumetric muscle loss injury. Journal of Biomedical Materials Research Part A 109, 695–712 (2021).

46 Huang, H. et al. The sustained PGE2 release matrix improves neovascularization and skeletal muscle regeneration in a hindlimb ischemia model. Journal of Nanobiotechnology 20, 95 (2022).

47 Turner, T. et al. Harnessing lipid signaling pathways to target specialized pro-angiogenic neutrophil subsets for regenerative immunotherapy. Science Advances 6, eaba7702 (2020).

48 Ley, K. et al. Neutrophils: New insights and open questions. Science immunology 3, eaat4579 (2018).

49 Frangou, E., Vassilopoulos, D., Boletis, J. & Boumpas, D. T. An emerging role of neutrophils and NETosis in chronic inflammation and fibrosis in systemic lupus erythematosus (SLE) and ANCA-associated vasculitides (AAV): Implications for the pathogenesis and treatment. Autoimmunity reviews 18, 751–760 (2019).

50 Fernandez-Yague, M. A. et al. Analyzing immune response to engineered hydrogels by hierarchical clustering of inflammatory cell subsets. Science Advances 8, eabd8056 (2022).

51 Dalli, J. & Serhan, C. Macrophage proresolving mediators—the when and where. Microbiology spectrum 4, 10.1128/microbiolspec.mchd-0001-2014 (2016).

52 Wosczyna, M. N. & Rando, T. A. A muscle stem cell support group: coordinated cellular responses in muscle regeneration. Developmental cell 46, 135–143 (2018).

53 Kano, Y., Padilla, D., Hageman, K. S., Poole, D. C. & Musch, T. I. Downhill running: a model of exercise hyperemia in the rat spinotrapezius muscle. Journal of Applied Physiology 97, 1138–1142 (2004).

54 Terjung, R. L. & Engbretson, B. M. Blood flow to different rat skeletal muscle fiber type sections during isometric contractions in situ. Medicine and Science in Sports and Exercise 20, S124–130 (1988).

55 Mackie, B. G. & Terjung, R. L. Blood flow to different skeletal muscle fiber types during contraction. American Journal of Physiology-Heart and Circulatory Physiology 245, H265–H275 (1983).

56 Lee, L. C., Liong, C.-Y. & Jemain, A. A. Partial least squares-discriminant analysis (PLS-DA) for classification of high-dimensional (HD) data: a review of contemporary practice strategies and knowledge gaps. Analyst 143, 3526–3539 (2018).

57 Ruiz-Perez, D., Guan, H., Madhivanan, P., Mathee, K. & Narasimhan, G. So you think you can PLS-DA? BMC bioinformatics 21, 1–10 (2020).

58 Fabregat, A. et al. Reactome diagram viewer: data structures and strategies to boost performance. Bioinformatics 34, 1208–1214 (2018).

59 Schiaffino, S., Rossi, A. C., Smerdu, V., Leinwand, L. A. & Reggiani, C. Developmental myosins: expression patterns and functional significance. Skeletal muscle 5, 1–14 (2015).

60 Esfandiari, E. et al. Long-term symptoms and function after war-related lower limb amputation: A national cross-sectional study. Acta orthopaedica et traumatologica turcica 52, 348–351 (2018).

61 Stinner, D. J. et al. Military and Civilian Collaboration: The Power of Numbers. Military medicine 182 (2017).

62 Faraji, E. et al. Health concerns of veterans with high-level lower extremity amputations. Military Medical Research 5, 1–10 (2018).

63 Xu, Z.-Z. et al. Resolvins RvE1 and RvD1 attenuate inflammatory pain via central and peripheral actions. Nature medicine 16, 592–597 (2010).

64 Ji, R.-R., Xu, Z.-Z., Strichartz, G. & Serhan, C. N. Emerging roles of resolvins in the resolution of inflammation and pain. Trends in neurosciences 34, 599–609 (2011).

65 Bang, S. et al. Resolvin D1 attenuates activation of sensory transient receptor potential channels leading to multiple anti-nociception. British journal of pharmacology 161, 707–720 (2010).

66 Lima-Garcia, J., et al. The precursor of resolvin D series and aspirin-triggered resolvin D1 display anti-hyperalgesic properties in adjuvant-induced arthritis in rats. British journal of pharmacology 164, 278–293 (2011).

67 Corona, B. T. et al. The promotion of a functional fibrosis in skeletal muscle with volumetric muscle loss injury following the transplantation of muscle-ECM. Biomaterials 34, 3324–3335 (2013).

68 Garg, K., Ward, C. L., Rathbone, C. R. & Corona, B. T. Transplantation of devitalized muscle scaffolds is insufficient for appreciable de novo muscle fiber regeneration after volumetric muscle loss injury. Cell and tissue research 358, 857–873 (2014).

69 Greising, S. M. et al. Unwavering pathobiology of volumetric muscle loss injury. Scientific reports 7, 13179 (2017).

70 Quarta, M. et al. Bioengineered constructs combined with exercise enhance stem cell-mediated treatment of volumetric muscle loss. Nature communications 8, 15613 (2017).

71 Gilbert-Honick, J. & Grayson, W. Vascularized and innervated skeletal muscle tissue engineering. Advanced healthcare materials 9, 1900626 (2020).

72 Greising, S. M., Corona, B. T., McGann, C., Frankum, J. K. & Warren, G. L. Therapeutic approaches for volumetric muscle loss injury: a systematic review and meta-analysis. Tissue Engineering Part B: Reviews 25, 510–525 (2019).

73 Castor-Macias, J. A. et al. Maresin 1 repletion improves muscle regeneration after volumetric muscle loss. Elife 12, e86437 (2023).

74 Han, W. M. et al. Synthetic matrix enhances transplanted satellite cell engraftment in dystrophic and aged skeletal muscle with comorbid trauma. Science advances 4, eaar4008 (2018).

75 Sollberger, G., Tilley, D. O. & Zychlinsky, A. Neutrophil extracellular traps: the biology of chromatin externalization. Developmental cell 44, 542–553 (2018).

76 Serhan, C. N. Treating inflammation and infection in the 21st century: new hints from decoding resolution mediators and mechanisms. The FASEB Journal 31, 1273 (2017).

77 Pharmaceuticals, T. TP-317 for Treatment of IBD, <https://www.thetispharma.com/ibd>

78 Dravid, A. A., M Dhanabalan, K., Agarwal, S. & Agarwal, R. Resolvin D1 -loaded nanoliposomes promote M2 macrophage polarization and are effective in the treatment of osteoarthritis. Bioengineering & Translational Medicine 7, e10281 (2022).

79 Norling, L. V. et al. Cutting edge: humanized nano-proresolving medicines mimic inflammation-resolution and enhance wound healing. The Journal of Immunology 186, 5543–5547 (2011).

80 Levy, E. S. et al. Tissue factor targeting peptide enhances nanoparticle binding and delivery of a synthetic specialized pro-resolving lipid mediator to injured arteries. JVS-Vascular Science 4, 100126 (2023).

81 Lance, K. D. et al. Unidirectional and sustained delivery of the proresolving lipid mediator resolvin D1 from a biodegradable thin film device. Journal of Biomedical Materials Research Part A 105, 31–41 (2017).

