## Supplemental for "Improving Functional Muscle Regeneration in Volumetric Muscle Loss Injuries by Shifting the Balance of Inflammatory and Pro-Resolving Lipid Mediators"

**This file includes:**

Figs. S1 to S7, Tables S1-S3

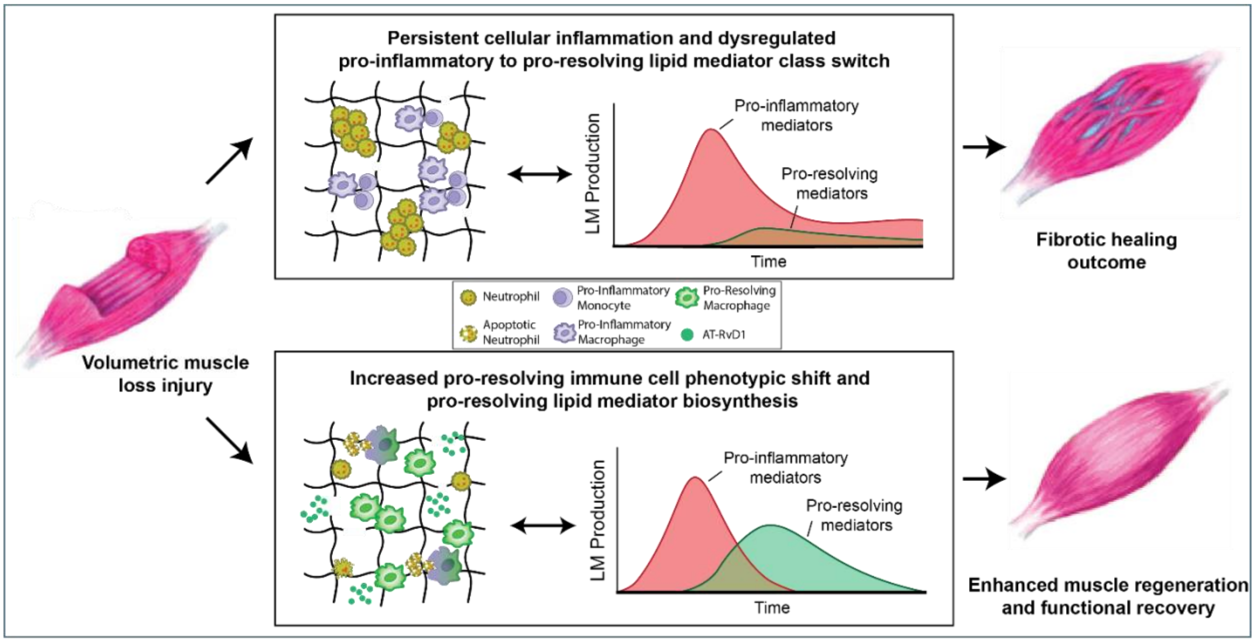


**Fig. S1 Graphical Abstract: A critically-sized VML injury was performed in a murine quadriceps model.** Unloaded (control) hydrogel treatment post-VML resulted in persistent inflammation and insufficient lipid mediator pro-inflammatory to pro-resolving class switching, leading to hallmark VML outcomes such as fibrosis rather than functional muscle recovery. In contrast, our localized, pro-resolving hydrogel releasing AT-RvD1 induced a shift towards a pro-resolving immune cell milieu and resulted in the increased biosynthesis of SPMs among various metabolic pathways which subsequently led to enhanced regeneration and improved functional recovery.

**Table S1: Antibodies used for cell phenotyping by flow cytometry.**

| Antibody | Fluorophore | Manufacturer |
|---|---|---|
| anti-CD29 | PE-Cy5 | Biolegend |
| anti-CD31 | APC | BioLegend |
| anti-Sca-1 | BV800 | Biolegend |
| anti-CXCR4 | PerCP-Cy5.5 | Biolegend |
| anti-Ly6C | BV510 | Biolegend |
| anti-CD206 | PE-Cy7 | Biolegend |
| anti-MerTK | PE | eBioscience |
| anti-CD64 | BV711 | Biolegend |
| anti-CD11b | PE-Cy5 | Biolegend |
| | BV421 | Biolegend |
| anti-Ly6G | APC-Cy7 | Biolegend |
| anti-CD49d | PE | Biolegend |
| anti-VEGFR1 | APC | Biolegend |
| Lineage Cocktail | APC | BD Pharmingen |
| anti-CXCR2 | PE-Cy7 | Biolegend |
| anti-CD47 | PerCP-Cy5.5 | Biolegend |
| anti-MPO | FITC | Biolegend |

**Table S2: Antibodies used for isolation of FAPs by FACS.**

| Antibody | Fluorophore | Manufacturer |
|---|---|---|
| anti-CD11b | APC | BioLegend |
| anti-CD31 | APC | BioLegend |
| anti-Ter119 | APC | BioLegend |
| anti-CD45 | APC | BioLegend |
| anti-Sca-1 | FITC | BioLegend |

**Table S3: Antibodies and dyes used for immunofluorescence staining of FAPs *in vitro*.**

| | Antibody/Stain | Fluorophore | Manufacturer | Cat. No. |
|---|---|---|---|---|
| **Primary** | Mouse anti-αSMA | None | Abcam | Ab7817 |
| | Rabbit anti-β1 integrin | None | ThermoFisher | PA5-78028 |
| | Hoechst 33342 | | Invitrogen | H3570 |
| | Phalloidin | i-Fluor 555 | Abcam | Ab176756 |
| **Secondary** | anti-mouse IgG | AlexaFluor 488 | ThermoFisher | A-11029 |
| | anti-mouse IgG | AlexaFluor 555 | ThermoFisher | A-21424 |
| | anti-mouse IgG | AlexaFluor 647 | ThermoFisher | A-21236 |
| | anti-rabbit IgG | AlexaFluor 488 | ThermoFisher | A-11034 |
| | anti-rabbit IgG | AlexaFluor 555 | ThermoFisher | A-21429 |
| | anti-rabbit IgG | AlexaFluor 647 | ThermoFisher | A-21245 |

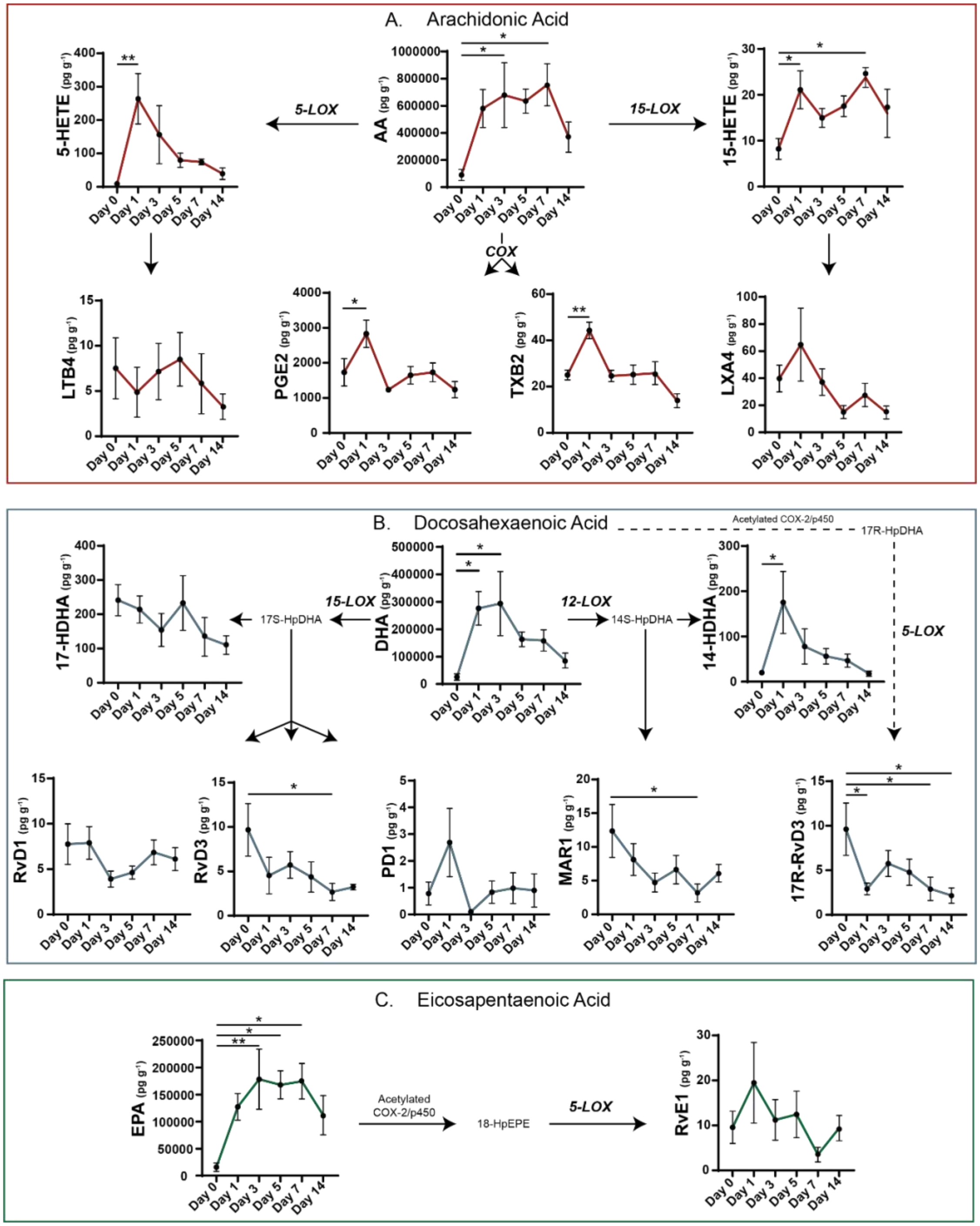


**Fig. S2: Concentration of downstream lipid mediators after VML injury demonstrates an inability of the tissue microenvironment to produce pro-resolving lipid mediators.** Quantification via LC-MS/MS of lipid mediators derived from AA (A), DHA (B), and EPA (C) of uninjured (day 0) quadriceps and days 1, 3, 5, 7 and 14 post critical VML injury. Total concentration of pro-inflammatory mediators and SPMs presented for uninjured (day 0) quadriceps muscle and indicated timepoints post-VML. Statistical analyses were performed using a one-way analysis of variance (ANOVA) with Dunnet's multiple comparisons; * $p < 0.05$ and ** $p < 0.01$; n=5 animals per group.

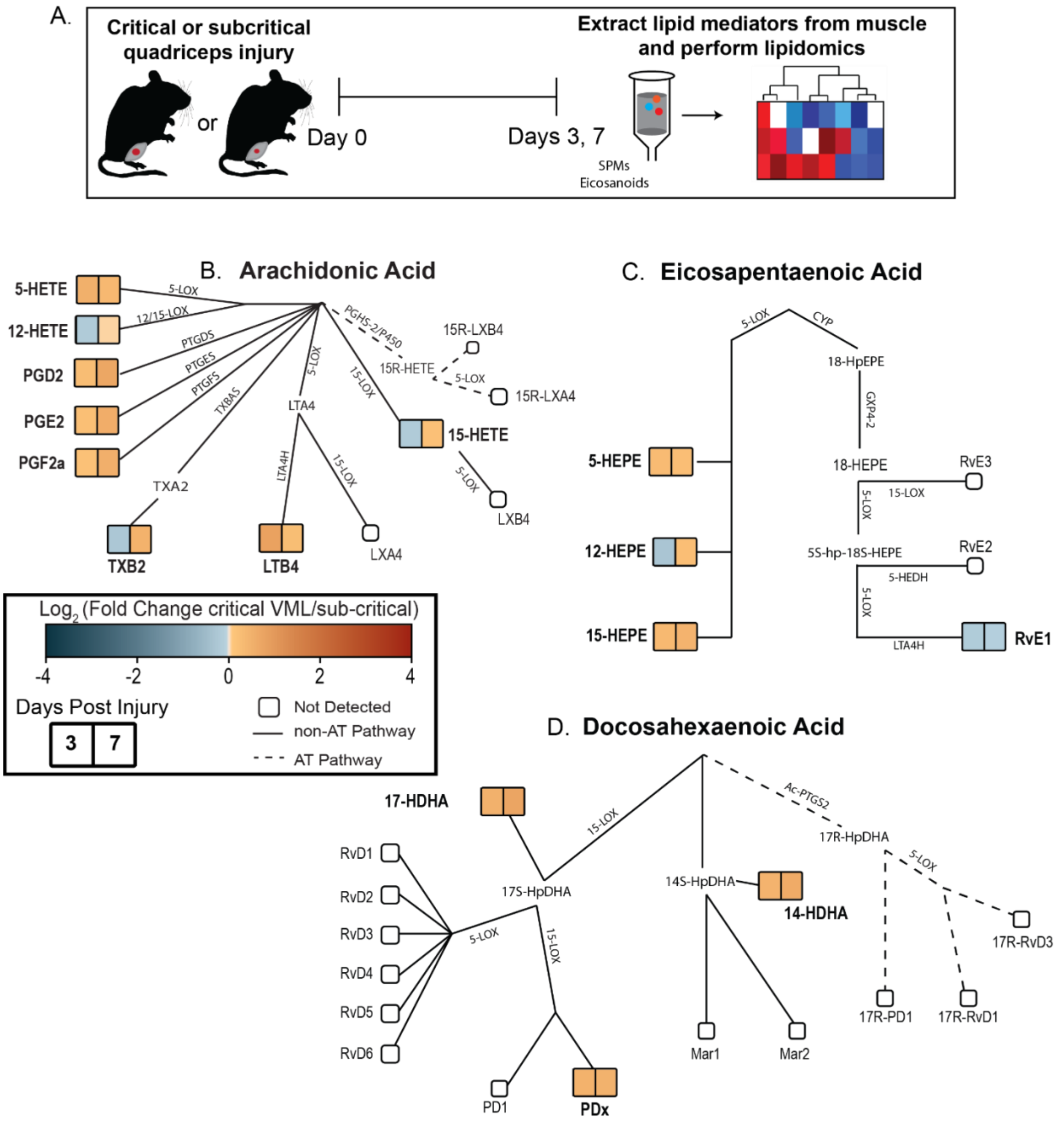


**Fig. S3: Targeted LC-MS/MS after VML reveals impaired production of specialized pro-resolving mediators relative to subcritical injury.** A subcritical injury (2mm biopsy) or critically-sized VML injury (3mm biopsy) was performed to the murine quadriceps, and resulting lipid mediators were analyzed via LC-MS/MS at day 3 and 7 post-VML (A). Heatmap rendering illustrating the fold change (critical VML/subcritical injury) of lipid mediators at 3- and 7-days post-injury overlaid onto the AA (B), EPA (C), and DHA (D) metabolic pathways. The metabolic pathway heatmap reveals that at days 3 and 7 post-VML the ability to produce pro-inflammatory eicosanoids (e.g. LTB4) is retained but the ability to produce SPMs is impaired. Heatmaps were created from the average concentration from n=5 animals per group.

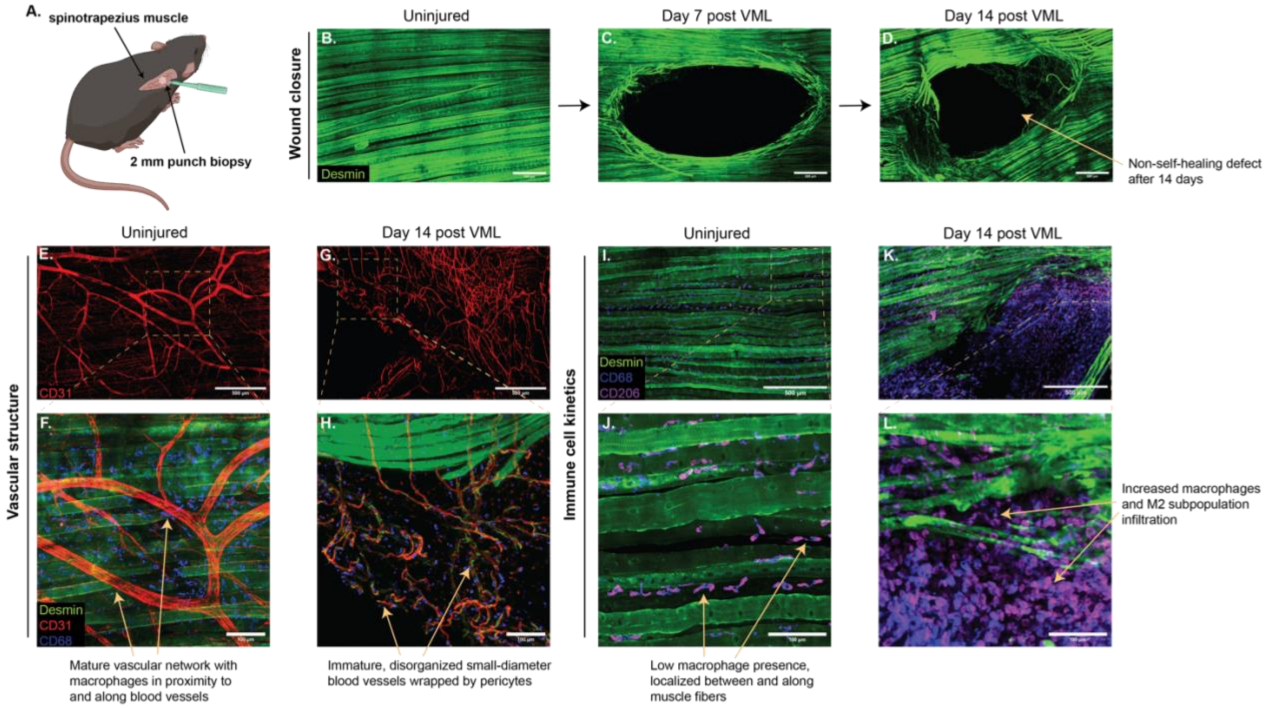


**Fig. S4: Spinotrapezius VML surgical model is established as a high-resolution platform for studying full-thickness tissue structure and cell interactions.** (A) A 2 mm-diameter full-thickness punch biopsy is performed through the right spinotrapezius muscle to create a VML defect. (B-D) Desmin (green) staining highlights the well-aligned muscle fiber structure in the uninjured spinotrapezius muscle (B) and demonstrates a reduction in wound size over time following VML, but also reveals the incapacity of the VML wound to self-heal after 14 days (C and D). (E-H) The vascular structure of the muscle is visualized by CD31 (red) staining. Uninjured spinotrapezius muscle possesses a mature vascular network with $CD68^+$ macrophages (blue) located near and along blood vessels (E and F). However, at day 14 post-VML, disrupted vascular structure is evident, with immature, disorganized small-diameter blood vessels enveloped by $Desmin^+$ pericytes (G and H). (I-L) The kinetics of macrophages and the M2 macrophage subpopulation are characterized by CD68 (blue) and CD206 (purple) dual staining. A small number of $CD68^+$ macrophages and $CD68^+CD206^+$ M2 macrophages are present in the uninjured spinotrapezius muscle (I and J), while 14 days after VML, there is increased infiltration and persistent presence of macrophages and the M2 subpopulation within the muscle (K and L).

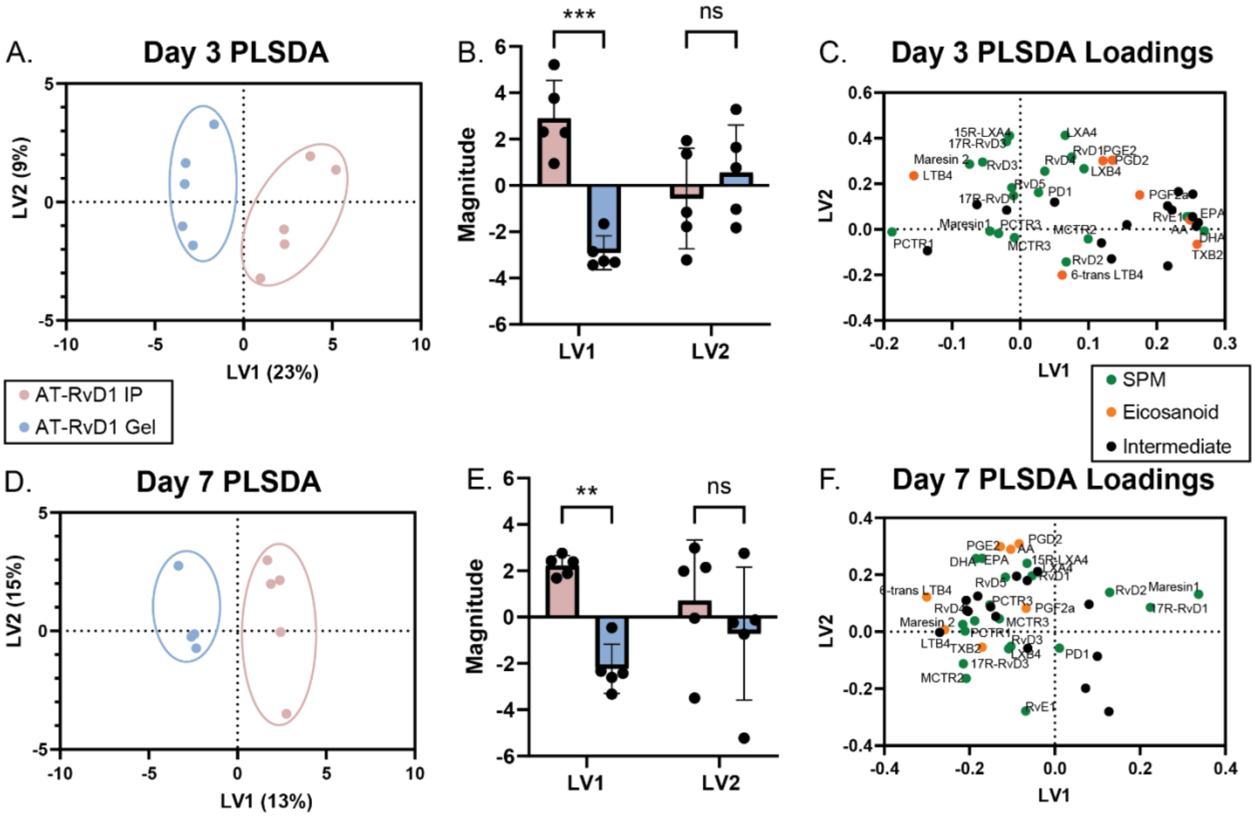


**Fig. S5: PLSDA reveals differential changes in overall quadriceps lipid mediator profile after local versus systemic administration of AT-RvD1.** Mice received either an empty VML defect with a single IP injection of AT-RvD1, or they were treated with local implantation of an AT-RvD1-loaded hydrogel. Separate PLSDA calculations were performed on the quadriceps lipid mediator profile at day 3 (A-C) or day 7 (D-F) post-injury. (A) and (D) Biplot of PLSDA embeddings on the first two latent variables at day 3 (A) and day 7 (D). Systemically- versus locally-treated groups are separated along the first latent variable at both day 3 (B) and day 7 (E). ** $p < 0.01$, *** $p < 0.001$. n=5 animals per group.

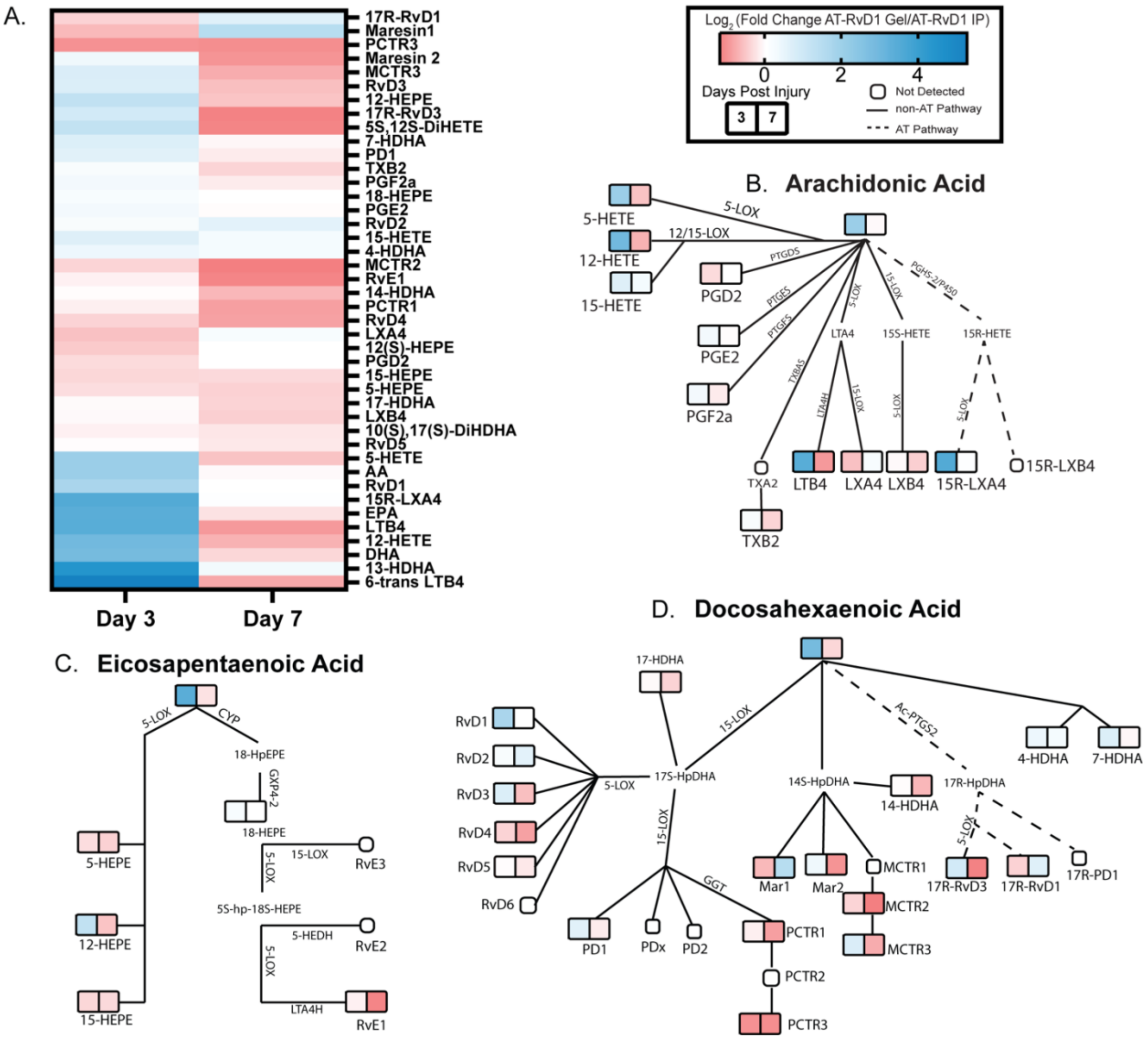


**Fig. S6: Distinct lipid mediator profiles occur in VML-injured quadriceps after systemic versus local AT-RvD1 treatment**. (A) The fold change of each lipid mediator's concentration between the local and systemic treatment groups were determined. Then the log2 of this value was calculated, represented in heatmap form. This heatmap is then overlaid onto the Arachidonic Acid (B), Eicosapentaenoic Acid (C) and Docosahexaenoic Acid (D) metabolic pathways. n=5 animals per group.

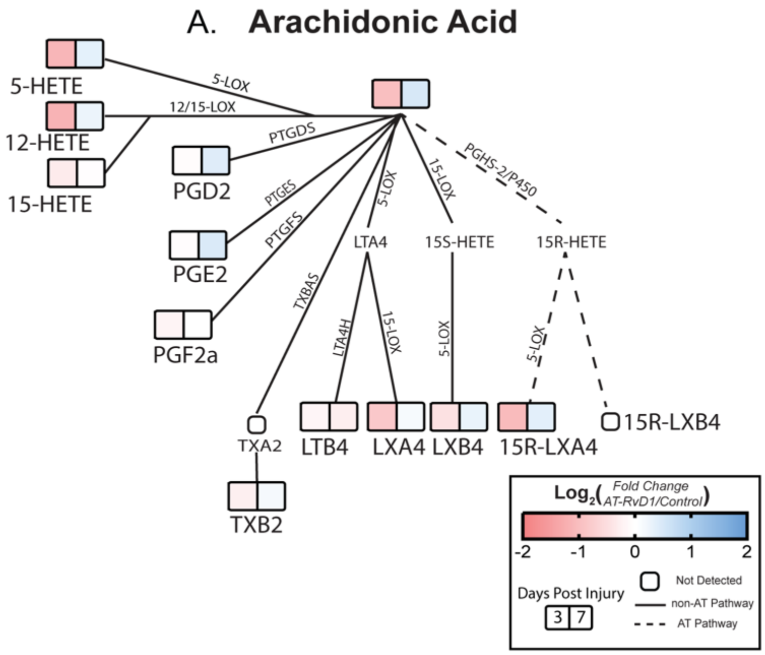


**Fig. S7:** The heatmap in Fig 6H overlaid onto wireframe diagrams of the Arachidonic acid (A) and Eicosapentaenoic acid (B) metabolic networks. Day 3 lipid mediator concentration is represented in the left square for each lipid mediator, while day 7 concentration is represented in the right square. n=5 animals per group.
